# Spatial distribution and cognitive impact of cerebrovascular risk-related white matter hyperintensities

**DOI:** 10.1101/2020.06.12.147934

**Authors:** Michele Veldsman, Petya Kindalova, Masud Husain, Ioannis Kosmidis, Thomas E. Nichols

## Abstract

**Objectives:** White matter hyperintensities (WMHs) are considered macroscale markers of cerebrovascular burden and are associated with increased risk of vascular cognitive impairment and dementia. However, the spatial location of WMHs has typically been considered in broad categories of periventricular versus deep white matter. The spatial distribution of WHMs associated with individual cerebrovascular risk factors (CVR), controlling for frequently comorbid risk factors, has not been systematically investigated at the population level in a healthy ageing cohort. Furthermore, there is an inconsistent relationship between total white matter hyperintensity load and cognition, which may be due to the confounding of several simultaneous risk factors in models based on smaller cohorts.

**Methods:** We examined trends in individual CVR factors on total WMH burden in 13,680 individuals (aged 45-80) using data from the UK Biobank. We estimated the spatial distribution of white matter hyperintensities associated with each risk factor and their contribution to explaining total WMH load using voxel-wise probit regression and univariate linear regression. Finally, we explored the impact of CVR-related WMHs on speed of processing using regression and mediation analysis.

**Results:** Contrary to the assumed dominance of hypertension as the biggest predictor of WMH burden, we show associations with a number of risk factors including diabetes, heavy smoking, APOE *ε*4/*ε*4 status and high waist-to-hip ratio of similar, or greater magnitude to hypertension. The spatial distribution of WMHs varied considerably with individual cerebrovascular risk factors. There were independent effects of visceral adiposity, as measured by waist-to-hip ratio, and carriage of the APOE *ε*4 allele in terms of the unique spatial distribution of CVR-related WMHs. Importantly, the relationship between total WMH load and speed of processing was mediated by waist-to-hip ratio suggesting cognitive consequences to WMHs associated with excessive visceral fat deposition.

**Conclusion:** Waist-to-hip ratio, diabetes, heavy smoking, hypercholesterolemia and homozygous APOE *ε*4 status are important risk factors, beyond hypertension, associated with WMH total burden and warrant careful control across ageing. The spatial distribution associated with different risk factors may provide important clues as to the pathogenesis and cognitive consequences of WMHs. High waist-to-hip ratio is a key risk factor associated with slowing in speed of processing. With global obesity levels rising, focused management of visceral adiposity may present a useful strategy for the mitigation of cognitive decline in ageing.

## 1. Introduction

White matter hyperintensities (WMHs) of presumed vascular origin (Wardlaw et al., 2013) are widely recognised as an indicator of poor brain health (Wardlaw et al., 2015). Age remains the strongest predictor for the presence of WMHs. However, the total burden of WMHs is higher in individuals with cerebrovascular risk (CVR) factors, like hypertension or hypercholesterolemia. WMHs triple the risk of stroke and double the risk of dementia suggesting they reflect pathological processes and are not simply a consequence of ageing (Debette and Markus, 2010). A number of studies have examined the relationship between different CVR factors and total WMH burden (Debette and Markus, 2010). Hypertension usually emerges as the dominant risk factor (Debette and Markus, 2010). Beyond this, it is less clear which risk factors are associated with the presence of WMHs in different regions of the brain, and with impaired cognition, when controlling for other risk factors. In other words, there is not sufficient evidence as to which risk factors make an independent contribution to WMH spatial distributions across the brain and associated cognitive impairment. This is important because it may change the focus of clinical management of risk factors beyond control of blood pressure.

Since the first visual scales attempting to quantify WMH burden (Fazekas et al., 1987), it has been recognised that WMHs disproportionately fall within periventricular (PV-WMHs) areas or in deep white matter regions (D-WMHs). Classification in this way has proven useful because deep and periventricular WMHs have different underlying microstructure, different associations with CVR factors (Griffanti et al., 2018) and potentially different relationships to cognition (Mortamais et al., 2013). Although pathology studies are relatively rare compared to imaging studies, there is some evidence of different pathological processes underlying WMHs in different regions (Wardlaw et al., 2015). PV-WMHs are associated with cerebral ischaemia and demyelination of adjacent fibre tracts as well as ependymal loss around the ventricles (Fazekas et al., 1993; Kim et al., 2008). PV-WMHs capping the ventricles are thought to be non-ischaemic in nature and reflect more generalised gliosis (Fazekas et al., 1998). In contrast, D-WMHs are thought to be of more ischaemic origin, the degree of confluence reflecting the degree of ischaemic damage with the most severe being marked loss of fibres and arteriolosclerosis (Fazekas et al., 1993). The spatial variability of WMHs associated with individual CVR factors has been investigated qualitatively using visual rating scales or broad region of interest approaches (De Leeuw et al., 2001; Strassburger et al., 1997; Van Dijk et al., 2004). There has been much less investigation of WMHs at a whole brain level, specifically looking at the probability of the presence of WMHs for a given risk factor, voxel-wise. Beyond the deep and periventricular classification, there may be important clues in the spatial distribution of WMHs that explain their pathogenesis and contribution to vascular cognitive impairment.

There are conflicting reports over whether hypertension, thought to be the strongest predictor of total WMH burden, is associated with D-WMHs specifically (Moroni et al., 2018; Strassburger et al., 1997) or more diffuse WMHs throughout the brain (Moroni et al., 2018; Wiseman et al., 2004). Diabetes presents a similarly confused picture in the literature. Some reports show no difference in total volume between diabetic patients and non-diabetic controls (De Bresser et al., 2018). One cross-sectional study showed increased D-WMH volume in diabetic patients as well as reduced blood flow. The reduced blood flow may explain the pathogenesis of diabetes related WMHs resulting from ischaemia in deep white matter (Abraham et al., 2016). Diabetes has also been associated with WMH load as a part of metabolic syndrome, showing a strong association with subcortical and periventricular WMHs (Abraham et al., 2016). However, neither body mass index (BMI), nor diabetes appeared to drive this relationship, instead hypertension was the predominant risk factor associated with WMH load. Hypertension frequently dwarfs the effects of the other CVR factors, and sample sizes are usually too low to examine the individual CVR factors - especially because of the high frequency of hypertension as comorbid with high BMI, diabetes and smoking.

The relationship between WMH load and smoking is also inconsistent across studies, but has been shown to be an independent risk factor, when controlling for age, in 1,814 participants of the Framingham Offspring cohort (Jeerakathil et al., 2004). BMI and visceral fat, either measured directly or indexed by waist-to-hip ratio (WHR), have also been associated with higher total WMH loads (Kim et al., 2017; Lampe et al., 2019b) and shown a preference for deep white matter, although this also varies by study (Griffanti et al., 2018; Lampe et al., 2019b). Finally, carriage of the apolipoprotein-E (APOE) *ε*4 allele is associated with higher total volume and higher accumulation of WMHs over time (Sudre et al., 2017), but also shows close interdependence with other CVR factors, such as hypertension (Salvado et al., 2019). Examining the independent contribution of the APOE *ε*4/*ε*4 genotype to the spatial distribution of WMHs is particularly difficult in studies with small cohorts, due to the relatively low prevalence of this allele in the general population (around 13%). Individual risk factors show some interaction with age, for example one large multi-centre stroke study in China showed high cholesterol to be a more important risk factor in older age (Ryu et al., 2014). Finally, sex appears to interact with some of the risk factors, such that the total WMH load is higher in females and the predictive risk factors different to males (Sachdev et al., 2009). Overall, what emerges from the literature is conflicting and complicated associations between individual risk factors and the presence of WMHs beyond those associated with age.

The literature to date has also shown a very mixed picture with regards to the relationship between total WMH load and cognition (Debette and Markus, 2010). Where a relationship has been observed, impairment to speed of processing and executive function are frequently associated with increasing total WMH load. A review of studies between 1990-2013 investigating the relationship between cognition and WMH load in the general population (Mortamais et al., 2013) found equivocal results. Five studies found a significant association between WMH load and global cognition and two reports failed to find an association (Mortamais et al., 2013). In studies that have investigated the spatial distribution of WMHs, cognitive decline was associated with periventricular WMHs (Godin et al., 2010; Prins et al., 2005). It is not clear whether particular risk factors increase the likelihood of cognitive decline associated with WMHs or whether it is just the total burden of global WMHs that is important.

The purpose of this study was to use population level imaging, demographic and lifestyle data from the UK Biobank to answer the following questions. Firstly, what is the cross-sectional relationship between total WMH load and age in the presence and absence of individual risk factors at the population level? This serves to clarify overall trends associated with the different risk factors and to highlight potential interactions between variables such as age and sex. Secondly, our main aim was to investigate whether the spatial distribution of WMHs, estimated voxel-wise across the whole brain, varied for individual CVR factors? Here, we were interested in the contribution of individual risk factors and the spatial distribution of WMHs whilst controlling for all other related CVR factors. We extend the literature in several ways, by demonstrating quantitative methods to estimate the spatial distribution, voxel-wise, and by taking advantage of a large sample to examine the unique effects of individual risk factors. Finally, we investigated the relationship between individual risk factors and speed of processing. We take advantage of a large dataset that enables systematic examination of individual risk factors, whilst controlling for other risks and a novel method to estimate the probability of CVR-associated WHMs at a voxel-wise level.

## 2. Methods

### 2.1. Participants

The study was conducted under Biobank application number 34077, and imaging data shared within the University of Oxford under application number 8107. UK Biobank participants gave written, informed consent for the study, which was approved by the Research Ethics Committee under application 11/NW/0382.

Participants were selected according to the flow chart in Figure 1, starting with 22,292 T1 images. Applying the published UK Biobank automated processing and quality control pipeline (Alfaro-Almagro et al., 2018), 1,985 T1 images were classified as non-usable (for a full list of T1 image imperfections see Table 3, Alfaro-Almagro et al. (2018)). Following the T2 FLAIR pipeline (Alfaro-Almagro et al., 2018), further 1,219 participants with T2 FLAIR images missing (incidents during acquisition, protocol changes) or non-usable (quality control including problems with the acquisition or with the registration to T1) were excluded, i.e. 20,188 participants with available WMH segmented brain images. We excluded data from individuals (590 total) with a current diagnosis or history of neurological or neuropsychiatric disease based on self-reported, non-cancer illness during a verbal interview with a trained nurse. Excluded diagnostic categories were traumatic brain injury, transient ischaemic attack, stroke, haematoma, infection of the nervous system, brain abscess, haemorrhage or skull fracture, encephalitis, meningitis, amyotrophic lateral sclerosis, multiple sclerosis, Parkinson’s disease, Alzheimer’s disease (AD), epilepsy or alcohol or drug dependency (see Table A.1 for a list of excluded participants by condition). Given known differences in cardiovascular disease and CVR factors between ethnicities (Benjamin et al., 2017; Howard, 2013), and the low proportion of non-white individuals in the Biobank cohort (Fry et al., 2017), we elected to exclude individuals who self-declared as non-white ethnicity (Figure 1). Participants with missing data and one individual with unusually high WMH load (about 12,000 voxels affected by WMHs) were also excluded. As a result, the final dataset for the analysis consisted of 13,680 individuals.

**Figure 1:**
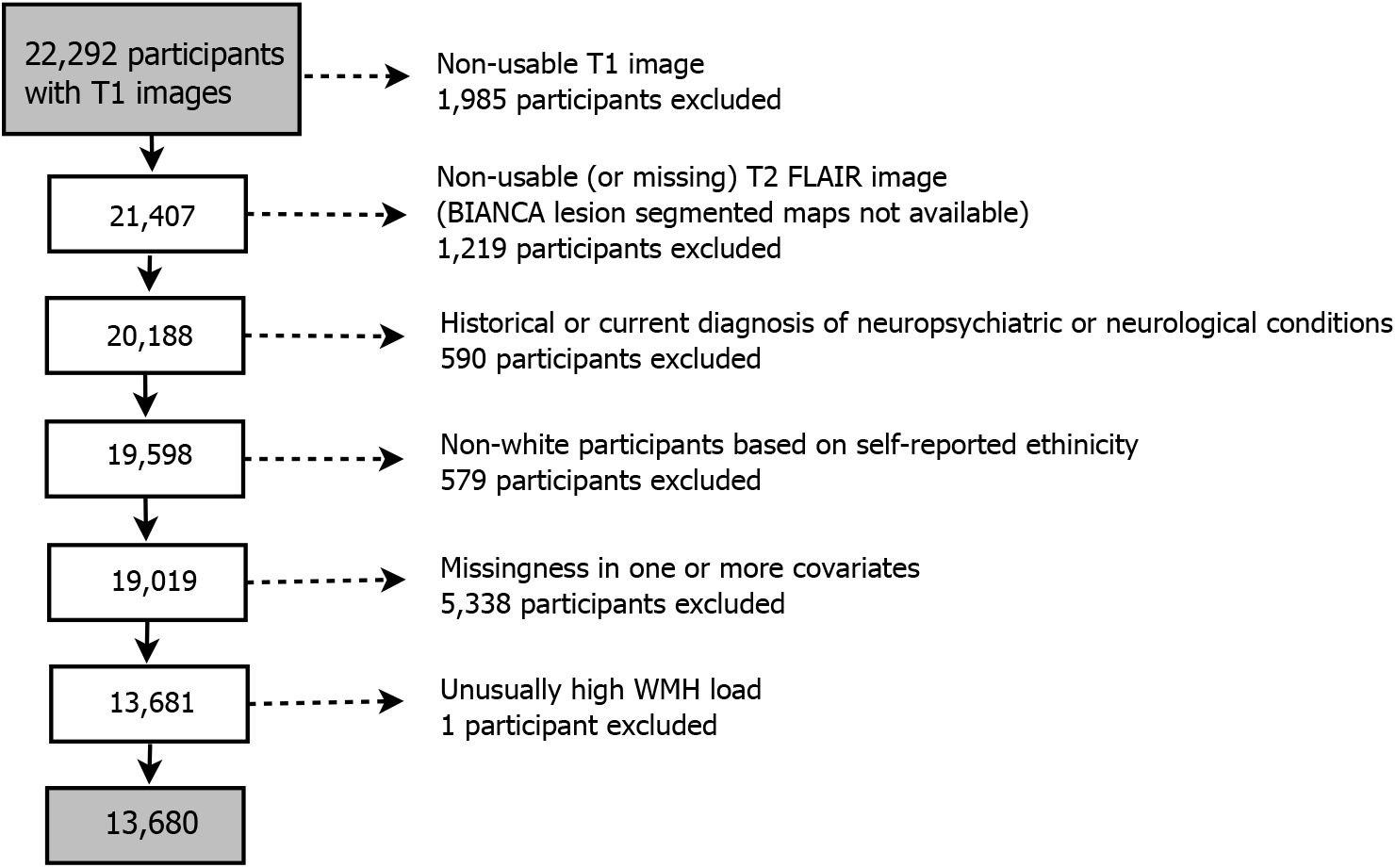
Diagram demonstrating the flow of gradually refining participants starting from all UK Biobank participants with available T1 structural brain images. Most common neuropsychiatric/neurological conditions in decreasing number of participants: stroke (185), transient ischaemic attack (115), epilepsy (77), etc. (see Table A.1 for a full list). Missingness by risk factor: hypertension risk (2,075), hypercholesterolemia (211), diabetes (165), smoking in pack years (3,146), waist-to-hip ratio (413), and APOE-*ε* status (514). Some individuals have more than one of the variables missing; for characteristics of individuals excluded due to missingness, see Table A.2.

### 2.2. Cerebrovascular risk factors

We investigated the main CVR factors known to be associated with the presence of WMHs, including hypertension, hypercholesterolemia, smoking, diabetes, waist-to-hip ratio and APOE-*ε* polymorphism status. Using categorical variables for each risk factor (described below), we also calculated a cumulative CVR score based on the sum of these variables to examine the impact of multiple risk factors on WMH distribution and its relationship to cognition. We did not use an established cardiovascular risk score, such as the Framingham Risk Score (Lloyd-Jones et al., 2004), for two reasons. Firstly, the established scores typically require continuous measures, such as high density lipoprotein levels, which were not available in UK Biobank at the time of analysis. Secondly, the established scores are typically used to calculate the future risk of a cardiovascular event, such as coronary heart disease. Here, we were interested in the cumulative effects of multiple CVR factors on WMH spatial distribution as well as on cognitive impairment, not on the risk of a specific cerebrovascular event.

#### Hypertension

Blood pressure (BP) was measured using a digital BP monitor (Omron), or a manual sphygmomanometer when the digital monitor was not available. Two readings were taken moments apart and we used the average of these two readings. To increase the reliability of our indicator for hypertension, we included self-reported medication for BP as an additional indicator of high BP. Therefore, our indicator variable ‘hypertension risk’ had value 1 for anyone (i) with average BP measuring over 140/90mmHg (Boffa et al., 2019) and/or (ii) on medication for high BP; otherwise the indicator had value 0.

#### Hypercholesterolemia

We used medication for cholesterol as an indicator for diagnosed hypercholesterolemia. Participants responded to the question “Do you take any of the following medications?” with the option of “cholesterol lowering medication” as part of a questionnaire presented on touch screen tablets. For those who selected “cholesterol lowering medication”, our indicator variable for hypercholesterolemia was assigned a value of 1, and 0 for those who answered this question but did not select this option.

#### Diabetes

Diabetes diagnosis was determined from responses to the question “Has a doctor ever told you that you have diabetes?”. This question was part of a questionnaire presented on touch screen tablets. Based on the answer to the question, the indicator variable for diabetes was 1 (answer ‘yes’) or 0 (answer ‘no’).

#### Smoking

Current and past smokers, and non-smokers, were divided into groups according to their pack year history. Pack years was calculated as the daily number of cigarettes divided by pack size (20) and multiplied by the number of years smoking. The number of years smoking was the age at stopping smoking, or the age at testing for current smokers, minus the age at which smoking was started. Pack years was appropriately adjusted for those who reported giving up smoking for more than six months. We grouped participants into non-smokers (less than or equal to 10 pack years), smokers (more than 10 and less than or equal to 50 pack years), heavy smokers (more than 50 pack years) (Lubin et al., 2016), which resulted in a discrete ‘smoking score’ covariate with three levels: non-smoker (0), smoker (1) and heavy smoker (2).

#### Waist-to-hip ratio

Waist circumference is a measure of visceral and subcutaneous fat, while hip circumference is thought to represent subcutaneous fat only. The ratio therefore represents an elevated proportion of intra-abdominal fat (Shuster et al., 2011). Waist and hip circumferences were manually measured in centimeters and used to calculate the WHR. The threshold for high WHR was set according to the WHO guidelines, set for each sex (World Health Organization, 2008) (0.9 for males and 0.85 for females). In the subsequent analysis we used WHR either as a continuous covariate, or as an indicator of high WHR (1) or not (0).

#### APOE-ε status

Carriage of the APOE *ε*4 allele only (not carriage of *ε*3/*ε*3 or *ε*2/*ε*2 or *ε*2/*ε*3) was considered a cerebrovascular risk factor based on a substantial body of research for APOE *ε*4 as a risk factor for cardiovascular disease (McCarron et al., 1999) and sporadic dementia. The genotyping pipeline is described in full here (Bycroft et al., 2018). Based on the number of *ε*4 alleles, a discrete APOE-*ε* status covariate was created as 0 (no carrier of *ε*4 allele), 1 (heterozygous, i.e. *ε*3/*ε*4) and 2 (homozygous, i.e. *ε*4/*ε*4).

#### CVR score

The CVR score was created as the sum of the six categorical variables representing the six risk factors described above: hypertension risk (0/1), hypercholesterolemia (0/1), diabetes (0/1), smoking score (0/1/2), WHR (0/1), and APOE-*ε* status (0/1/2). The resulting composite score was on a scale 0–8 and the higher the score, the higher the cerebrovascular burden of an individual. The UK Biobank data used to obtain the score are listed in Table A.3.

### 2.3. Cognitive testing

Data from the reaction time task, thought to be a sensitive index of speed of processing, was used in the analysis (Fawns-Ritchie and Deary, 2020). We used speed of processing as a cognitive variable because it has most consistently shown a relationship with WMH load, it is normally distributed and it has considerably less missing data than some of the other cognitive test variables available in UK Biobank. The task was administered by touch screen at the same session as the MRI scan. The UK Biobank reaction time task was a variant of the ‘Snap’ card game, in which participants react to the presence of a pair of matching cards over 12 rounds of the game. Mean reaction time was recorded and trials with responses below 50ms or above 2000ms were excluded.

Education is considered an important confounding variable when modelling cognitive function (Evans et al., 1993; Whalley et al., 2004). Participants responded to the question “Which of the following qualifications do you have?” as part of the questionnaire presented on touch screen tablets. A continuous variable “years of education” was defined according to the ISCED categories (Cheesman et al., 2020; Lee et al., 2018); for details see Table A.3.

Missingness in the cognitive task variable resulted in 923 exclusions with further 27 exclusions due to missingness in the education variable. Those participants were excluded only when the statistical analysis included reaction time as a variable, otherwise 13,680 individuals were used.

### 2.4. MRI data

Volunteers were scanned on Siemens Skyra 3T scanners with 32 channel head coils. We used the T2-weighted fluid attenuated inversion recovery (FLAIR, 1.05 × 1 × 1mm resolution) and the T1-weighted, 3D magnetization-prepared rapid gradient echo (MPRAGE, 1×1×1mm resolution, T1=880ms, TR=2000ms, matrix=208×256×256) sequence as part of the longer imaging protocol^1^. The published UK Biobank pipeline (Alfaro-Almagro et al., 2018) details the spatial normalisation procedure of the T1 image to MNI 152 space. Briefly, after gradient distortion correction and reduction of the field of view (FOV) to remove non-brain space, FNIRT (Anderson et al., 2007) was used for non-linear registration to 1mm resolution MNI 152 space. FNIRT parameters were optimised for best performance on the UK Biobank’s T1 image resolution and contrast.; the FNIRT configuration file used as part of the UK Biobank pipeline is available online at https://git.fmrib.ox.ac.uk/falmagro/UK_biobank_pipeline_v_1/-/blob/master/bb_data/bb_fnirt.cnf. All three of the above steps are combined into a single non-linear and reversible transformation. Note that participants with large ventricles were excluded from the dataset as part of the UK Biobank quality control pipeline. After gradient distortion correction, the T2 FLAIR image in native space was rigid-body transformed using FLIRT (Jenkinson et al., 2002) to register to T1 space. We excluded individuals with non-usable or missing T1 or T2 FLAIR images, see Section 2.1 and Figure 1.

The Brain Intensity Abnormality Classification Algorithm (BIANCA) (Griffanti et al., 2016) was used to segment WMHs. BIANCA is an automated method for WMH segmentation based on voxel intensity and distance to the ventricles. BIANCA’s segmentation has been compared to segmentation of WMHs from two different cohorts, with different MRI sequence parameters, as well as in patient populations (a vascular and a neurodegenerative cohort). Correlations in total extracted WMH load, spatial overlap and visual rating scales has shown BIANCA to be a valid alternative to manual segmentation (Griffanti et al., 2016). When applied to the UK Biobank imaging data, it produced an output image in subject space which represented the probability per voxel of being a WMH; as part of the segmentation, it was then thresholded at 0.8 to give a binary WMH mask. The threshold of 0.8 was the optimised tuning parameter to minimise prediction error in native space when compared to manually segmented lesion masks of 12 UK Biobank individuals. We applied the estimated spatial normalisation parameters to the WMH maps; specifically, the transformation parameters for the T2-weighted FLAIR and non-linear warping were used (Andersson et al., 2007), and then we thresholded the warped WMH maps (the warping process used trilinear interpolation that introduced non-binary values) at 0.5 to get binary WMH maps in MNI space. The 0.5 threshold was used as a neutral value to preference neither enlargement or shrinkage of total lesion volume. Having a binary WMH map per participant, we estimated the WMH load as the number of WMH-affected voxels.

All preprocessing steps were performed using the FSL software^2^. Note that voxel size of 2mm^3^ was chosen for computational reasons and this implied a standard brain mask of dimension 91 × 109 × 91 voxels. Binary WMH maps and WMH load (unit of measurement was 2mm^3^) were available for 13,680 participants.

### 2.5. Statistical analysis

We chose to do complete cases analysis, rather than impute missing data, and therefore excluded 5,338 individuals with missing data in one or more CVRs (see Figure 1 for details). All the exclusions as described in Section 2.1 led to a final dataset for the analysis of 13,680 participants.

#### Univariate analysis

From each participant’s WMH binary mask, where one indicates the presence of a WMH and zero the absence, respectively, we used the subject-specific summary measure log-transformed WMH load as the response variable in our first modelling step. Given the highly right-skewed distribution of WMH load across the population, we log-transformed WMH load to enable the use of standard least squares linear models.

As an exploratory step of whether and how aging related to log(WMH load) in individuals grouped by presence or absence of cerebrovascular risk factors (or by sex), we used locally estimated scatterplot smoothing (loess) (Cleveland et al., 2017) as implemented in `R` package `stats`, function `loess.smooth()`. Loess is a non-parametric local averaging method, which uses weighted regression inside windows with a fixed number of points. To determine the window, we fixed the span parameter to 20%, which means that the horizontal window surrounding a target observation contains 20% of its nearest neighbours. Then, a weighted polynomial was fitted to the data within the window and the predicted response at the target point was the fitted value. The fitted smooth curve provided a graphical overview of underlying patterns in the dataset. We used the resulting fitted curves to explore (i) the linear age effect assumption, (ii) the need for an age by sex interaction, and (iii) the effect of risk factors on log(WMH load) and whether it varied across age.

Multiple linear regression was used to formally assess the dependence of log(WMH load) on the cerebrovascular risk factors, while controlling for known confounders (age, sex, head size). We controlled for head size which is a recognised confounding variable in MRI studies generally and in UK Biobank specifically (Alfaro-Almagro et al., 2020). Because head size correlates with sex, spurious correlations can arise between sex and MRI variables if head size is not controlled for. A recent paper on deconfounding UK Biobank MRI data (Alfaro-Almagro et al., 2020), recommends the minimal set of confounding variables includes age, sex, age-sex interaction and head size scaling (Section 2.4.1). We also explored the inclusion of an age-sex interaction term to the models as one of the confounds.

With *N* subjects and *Y*_*i*_ the random variable representing the log(WMH load) for each subject *i* (*i* = 1, …, *N*), suppose all *Y*_*i*_’s are independent and Normally distributed with means 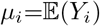 and variance *σ*^2^, which is the same across subjects. Then a normal linear model can be written as

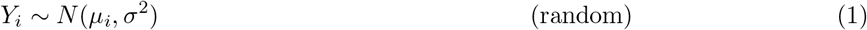

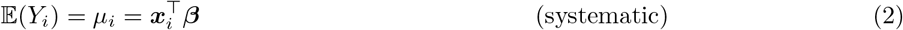

where *β* is an *P*-vector of parameters, and *x*_*i*_ denotes the *P*-vector of subject-specific covariates for subject *i* and *X* is the full rank design matrix with rows ***x***_1_, …, *x*_*N*_.

To assess the importance of each explanatory variable, maximum likelihood estimates (MLEs) of the regression coefficients were explored along with 95% confidence intervals (CIs) and p-values (significance level 0.05). Note that for the discrete explanatory variables (such as sex, hypertension risk, smoking score, etc.), level 0 of the factor variable was fitted to be the baseline, e.g. the estimated regression coefficient 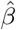 for hypertension risk estimated the effect of hypertension risk 1 on the outcome in comparison to the effect of hypertension risk 0. If the factor variable had more than 2 levels, here smoking score and APOE-*ε* status, a dummy variable was created for each of the two contrasts with category 0 used as baseline - category 2 compared to category 0 and category 1 compared to category 0 - so two regression coefficients were estimated for the two contrasts. We used adjusted *R*^2^ 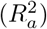 as a measure of goodness of fit, which is interpreted as the percentage of variance explained. We also computed partial 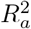, which measures the additional variation explained by each explanatory variable, after adjustment for the other predictors. We fitted various multiple regression models aiming to (i) outline the risk factors which were significant predictors of log(WMH load), and (ii) to determine the models used for the spatial voxel-wise analysis.

#### Voxel-wise analysis

Voxel-wise analysis was employed to assess how different contributors to the cerebrovascular burden related to the spatial distribution of WMHs.

Mass-univariate voxel-wise modelling of WMH masks requires a generalized linear model (GLM), e.g. logistic regression or probit regression, to account for the binary nature of the WMH masks. Since we now want to model WMH probability at each voxel, let *Y*_*i*_(*s*_*j*_) denote a Bernoulli random variable with probability of success *p*_*i*_(*s*_*j*_), where *Y*_*i*_(*s*_*j*_) represents the presence (*Y*_*i*_(*s*_*j*_)=1) or absence of a WMH for subject *i, i* = 1, …, *N* at voxel *s*_*j*_ (*j* = 1, … , *M*). Assume *Y*_1_(*s*_*j*_), … , *Y*_*N*_ (*s*_*j*_) are independent random variables across subjects *i* and voxels *s*_*j*_. In contrast to the normal linear model, every GLM has a link function *g*, which is a monotonic function that relates the expectation of the random outcome to the systematic component. Note that the link function is the identity link function for the normal linear model in Equations (1, 2). The GLM can be written as

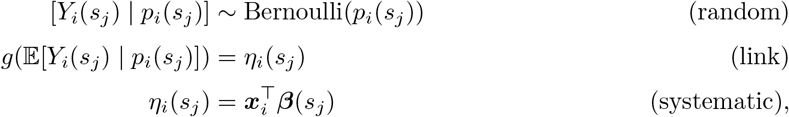

where ****β****(*s*_*j*_) is a *P*-vector of parameters at each voxel *s*_*j*_.

For this analysis, we have chosen probit link^3^ Φ^−1^, where Φ indicates the standard normal cumulative distribution function, so the model can be written as probit regression

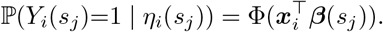

At each voxel *s*_*j*_, we obtain the MLEs 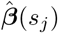 through iterative optimization, such as iteratively reweighted least squares (IWLS) (Green, 1984). The MLE is the default choice of an estimator due to its optimal asymptotic properties. However, in finite samples, the MLE may demonstrate significant bias and large variance. Furthermore, in binary response models, there is positive probability for the MLE to have at least one infinite component, which results in issues with common inference procedures, such as Wald tests and Wald-type CIs. Such infinite estimates result when the data are separated (Albert and Anderson, 1984), that is a covariate or a combination of covariates perfectly separates outcome measurements (e.g. female participants in the dataset having a WMH at a particular voxel where no male does). If the MLE is infinite, inference becomes impossible and test statistics are unstable.

Logistic or probit regression has been often used in the voxel-wise brain WMH mapping literature (Lampe et al., 2019b; Rostrup et al., 2012), but potential convergence and bias problems have not been discussed to our knowledge. To address both limitations-infinite and biased MLEs - we propose the use of test statistics (standardized coefficients) based on mean bias-reduced (BR) estimates 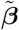. The bias-correction approach, which we focused on in the work, was first introduced in Firth (1993) for logistic regression and then further developed for exponential family GLMs (Kosmidis and Firth, 2009; Kosmidis et al., 2020). This method guarantees finite estimates when total separation is observed and it ensures estimates with asymptotically smaller bias than what the MLE has. Bias reduction is achieved by subtracting the first-order bias in each iteration of the optimization; see Kosmidis and Firth (2009) for the exact form of the adjustments. To obtain mean BR estimates, we use the `R` package brglm2 (Kosmidis et al., 2020), which adds additional functionalities to the `R` function `glm()`. Note that as in the univariate analysis, we chose level 0 as the baseline for factor variables, i.e. one regression coefficient was estimated voxel-wise for binary explanatory variables and two for discrete variables with three levels (level 2 vs baseline (level 0) and level 1 vs baseline), respectively.

To determine the size of the effect of each explanatory variable, we explored test statistics (standardized coefficients or z-scores) based on mean BR estimates 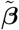 and their associated p-values. While the mean BR estimates ensure better performance when there are few WMH at a voxel, like other authors we excluded voxels when the WMH count fell too low; for example, Rostrup et al. (2012) and Lampe et al. (2019b) required at least 5 participants. Due to the large sample size, we chose 4 as our threshold, i.e. we only considered voxels where 4 or more participants had a WMH (see Appendix B for more details).

After computing the p-values across the brain, we corrected for multiple testing. Threshold-free cluster enhancement (TFCE) (Smith and Nichols, 2009) is often used in the literature, but due to the UK Biobank sample size of 13,680, TFCE would be computationally expensive to perform (c.f. 605 for Rostrup et al. (2012) and 1,825 for Lampe et al. (2019b), where authors employed the TFCE approach). Instead, we used false discovery rate (FDR) correction (Benjamini and Hochberg, 1995), i.e. we controlled the expected proportion of falsely rejected hypotheses. We favoured FDR over the most common family-wise error rate correction method, Bonferroni correction, since Bonferroni is known to be quite conservative when the comparisons are not independent, which is the case for spatially dependent WMH maps (Genovese et al., 2002; Rorden and Karnath, 2004). The `R` function `p.adjust()` (package `stats`) was used to correct the p-values, using the *α*_FDR=0.05_ significance level. We visually inspected axial slices of the the standardized coefficients for ‘significant’ voxels (voxels where an explanatory variable was a significant predictor of WMH risk) to gain understanding of the localized effect of cerebrovascular risk factors and how they complemented each other. We also inspected the total number of significant coefficients across the brain per predictor across a variety of models as a measure of the spread of the effect throughout white matter.

#### Mediation analysis

The univariate analysis framework was also employed to explore the association between speed of processing (reaction time) and log(WMH load) (or CVR score). To better understand the underlying dependencies, we also performed mediation analysis to investigate the hypothesis that the effect of WMH load on speed of processing (cognitive task) was fully or partially explained through a given CVR (mediator); see Figure A.3. The `R` package `mediation` (Tingley et al., 2014) was used for the estimation and suitable models (GLMs, LMs) were used for the mediator and outcome models (some of the mediators are discrete variables, which necessitates the use of GLMs). All models were controlled for age, sex, age by sex interaction, head size and years of education. Point estimates along with 95% percentile CIs and p-values were explored for the direct, indirect and total effects (non-parametric bootstrap, 10,000 resamples). We were mostly interested whether the indirect effect was significant, i.e. whether there was significant mediation effect, and if so, what was the proportion mediated (percentage of total effect) of WMH load on speed of processing operating (partially or fully) through the CVR factors.

## 3. Results

We analysed data from 13,680 individuals from the UK Biobank (mean age 62.9 ± 7.4 years, 7,236 female). Summary descriptive statistics of the UK Biobank sample characteristics are included in Table 1. The UK Biobank variables used in the current work are described in Table A.3.

**Table 1:** Characteristics of UK Biobank dataset of 13,680 participants (*12,730 participants for the cognition variable reaction time and education due to missingness). SD: standard deviation; WHR: waist-to-hip ratio; APOE: apolypoprotein-E; CVR: cerebrovascular risk; WMH: white matter hyperintensity.

| Characteristics | Levels (N) | Mean (SD) | Median (range) |
| --- | --- | --- | --- |
| Age (years) | — | 62.9 (7.4) | 63.5 (45.1; 80.7) |
| Sex | Men (6,444), Women (7,236) | — | — |
| Head size | — | 1.3 (0.1) | 1.3 (0.9; 1.8) |
| Hypertension risk | 0 (7,272), 1 (6,408) | — | — |
| Hypercholesterolemia | 0 (10,899), 1 (2,781) | — | — |
| Diabetes | 0 (13,017), 1 (663) | — | — |
| Smoking (score/pack years) | 0 (11,291), 1 (2,238), 2 (151) | 4.9 (11.4) | 0 (0; 141) |
| WHR (indicator/continuous) | 0 (7,251), 1 (6,429) | 0.9 (0.1) | 0.9 (0.6; 1.2) |
| APOE-ε status | 0 (10,226), 1 (3,150), 2 (304) | — | — |
| CVR score | 0 (2,596), 1 (4,204), 2 (3,770),<br>3 (1,947), 4 (869), 5 (252),<br>6 (36), 7 (6), 8 (0) | 1.7 (1.2) | 2 (0; 7) |
| WMH load (2mm<sup>3</sup> voxels) | — | 528.6 (667.7) | 310.0 (3; 8,228) |
| Reaction time* (milliseconds) | — | 585.4 (104.9) | 569 (272; 1,559) |
| Years of education | 7 (780), 10 (1,679), 13 (757), 15 (1,481), 19 (2,027), 20 (6,006) | 16.7 (4.3) | 19 (7; 20) |

### 3.1. Age by sex interactions

Figure 2 suggests a positive linear relationship between age and CVR score as well as between age and log(WMH load). Males had overall higher cerebrovascular burden than females with no interaction with age (loess curves nearly parallel). Log(WMH load) increased with age and the different slope of the sex-specific fitted curves suggested a potential age by sex interaction. This was further confirmed through a highly significant interaction term (*p*<0.001) in a linear model of log(WMH load), adjusted for head size and total CVR burden (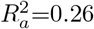, Table 2). To understand the effect of sex on the log(WMH load), we used the estimated regression coefficients for sex and age:sex interaction term (Model U.1) to check how the outcome variable changed. A female participant aged 75 years would be expected to have exp(−0.48 + 0.01×75)= exp(0.27)=1.31-fold higher WMH load than a male participant the same age. For a female participant aged 50, we get 1.02-fold difference in WMH load, respectively, which highlights the effect of the interaction term. We therefore adjusted all subsequent univariate models of log(WMH load) and voxel-wise models of WMH masks for an age by sex interaction, something often overlooked in analyses within the existing literature.

**Figure 2:**
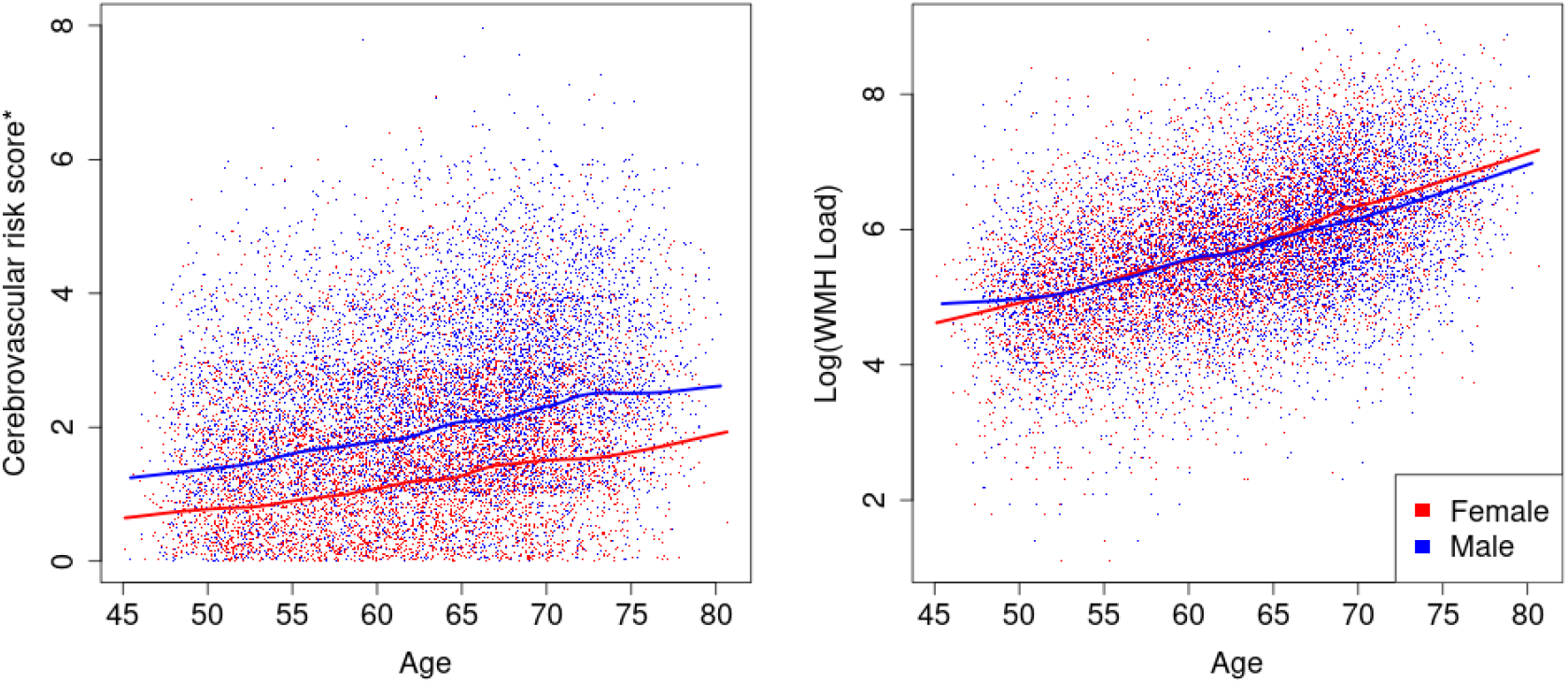
Sex-specific trends in CVR score (left) and in log(WMH load) (right) across age. Solid lines represent the loess-smoothed curve with a span of 20% and the points are the observed data points. Males have higher CVR burden than females across all ages and log(WMH load) increases across age for both sexes but potentially at different speed for males and females. *uniform noise U(0, 1) added to the CVR scores to disperse the values in the y-axis direction (left plot).

**Table 2:** Univariate regression summaries outlining the association between log(WMH load) and cerebrovascular risk factors (or composite score). For all CVRs, the presence of the risk has a strong positive effect on log(WMH load) when compared to its absence (discrete risk equals 0), e.g. participants who have high WHR are expected to have 0.22-fold higher log(WMH load), or exp(0.22) = 1.25-fold higher WMH load, than those who do not. Model U.1: all predictors in the model shown. Model U.2.1 - U.2.6: main effect of interest shown. All models adjusted for age, sex, head size, age-sex interaction and only one of the risk factors included in the 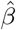 model. stands for the maximum likelihood estimate of the regression coefficient *β*; two regression coefficients for discrete variables with more than two levels (models U.2.4 and U.2.6): contrasting level 1 to level 0, and level 2 to level 0 (level 0 modeled as baseline).

| Model / Predictor (level) | Estimate $\hat{\beta}$ | 95% CI | p-value | $R_a^2$ / partial $R_a^2$ |
| --- | --- | --- | --- | --- |
| Model U.1 |  |  |  | **0.260** |
| Intercept | 2.19 | (1.93; 2.46) | <0.001 |  |
| Age | 0.06 | (0.05; 0.06) | <0.001 | 0.093 |
| Sex (Female) | -0.48 | (-0.74; -0.22) | <0.001 | 0.001 |
| Age:Sex (Female) | 0.01 | (0.01; 0.01) | <0.001 | 0.002 |
| Head size | -0.29 | (-0.45; -0.13) | <0.001 | 0.001 |
| CVR score | 0.14 | (0.13; 0.15) | <0.001 | 0.031 |
| Model U.2.1 |  |  |  | **0.249** |
| Hypertension risk (1) | 0.24 | (0.21; 0.28) | <0.001 | 0.017 |
| Model U.2.2 |  |  |  | **0.240** |
| Hypercholesteromia (1) | 0.17 | (0.13; 0.21) | <0.001 | 0.005 |
| Model U.2.3 |  |  |  | **0.240** |
| Diabetes (1) | 0.30 | (0.22; 0.37) | <0.001 | 0.005 |
| Model U.2.4 |  |  |  | **0.240** |
| Smoking score (1) | 0.15 | (0.11; 0.19) | <0.001 | 0.006 |
| Smoking score (2) | 0.45 | (0.30; 0.59) | <0.001 |  |
| Model U.2.5 |  |  |  | **0.245** |
| Waist-to-hip ratio (1) | 0.22 | (0.15; 0.25) | <0.001 | 0.012 |
| Model U.2.6 |  |  |  | **0.237** |
| APOE-ε status (1) | 0.05 | (0.01; 0.08) | 0.011 | 0.002 |
| APOE-ε status (2) | 0.23 | (0.13; 0.34) | <0.001 | |

**Table 3:** Percentage and number of significant voxels across predictors for joint model S.2 (all CVR factors) and for marginal models S.3.1 - S.3.6 (one of the CVR factors). Note that for factor variables with more than two levels (smoking score and APOE-*ε* status), level 0 is used as a baseline and we estimate the effects of level 1 and level 2 relative to baseline. All models include the same confounding variables age, sex, age-sex interaction and head size; Voxels with at least four individuals having a WMH explored (40,001 voxels in the brain mask) and 5% FDR correction applied, i.e. % FDR-corrected voxels is out of a total of 40,001 voxels. Columns 2 and 3 complementary to Figure 6(a) and 6(b), respectively.

| Predictor (level) | FDR-corrected voxels % (voxel count) |  |
| --- | --- | --- |
|  | Joint model (S.2) | Marginal models (S.3.1-S.3.6) |
| Hypertension risk (1) | 11.8% (4,705) | 13.4% (5,366) |
| Hypercholesterolemia (1) | 0.02% (10) | 1.6% (648) |
| Diabetes (1) | 1.5% (607) | 8.2% (3,270) |
| Smoking score (1) | 0.3% (133) | 1.6% (636) |
| Smoking score (2) | 1.1% (424) | 4.1% (1,633) |
| Waist-to-hip ratio (1) | 4.6% (1,841) | 10.7% (4,297) |
| APOE-ε status (1) | 0.0% (0) | 0.0% (0) |
| APOE-ε status (2) | 4.2% (1,708) | 3.9% (1,549) |

### 3.2. White matter hyperintensity load associated with individual risk factors

Next, we explored the relationship between log(WMH load) and individual CVR factors, including hypertension risk, hypercholesterolemia, diabetes, smoking, WHR, and APOE-*ε* status through fitting linear regressions (Table 2) and risk factor specific loess-smoothed curves (Figure 3). Age’s dominant effect was seen here: a 10-year age difference was associated with a exp(0.6)=1.82-fold difference in WMH load. All CVR factors had a marked positive effect on WMH load (Table 2), with hypertension risk and WHR having the highest partial *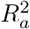* (1.7% and 1.2%), e.g. hypertension risk explained 1.7% of the remaining variability after adjusting for confounding. The inclusion of the CVR score as a predictor led to achieving the highest 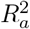 and its partial *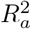* of 3.1% resembled the combined effect of the risk factors.

**Figure 3:**
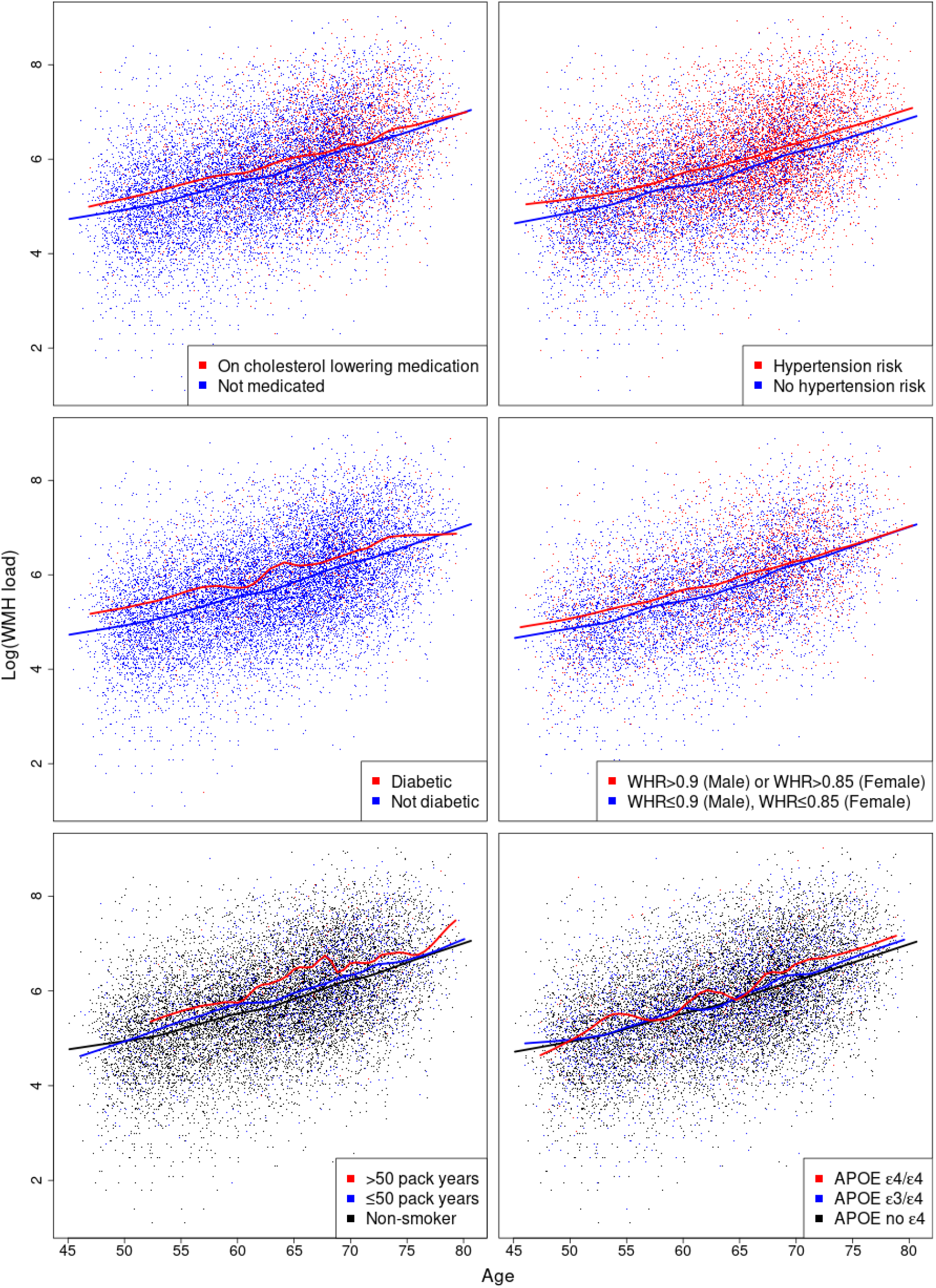
Cerebrovascular risk factor specific trends in log(WMH load) across age. Solid lines represent the loess-smoothed curve with a span of 20% and the points are the observed data points. The presence of any of the risk factors suggests higher log(WMH load). Crossing fitted curves would suggest a potential risk factor by age interaction and parallel line its absence, respectively. WMH: white matter hyperintensity; WHR: waist-to-hip ratio; APOE: apolypoprotein-E.

Hypertension risk was associated with 1.27-fold higher total WMH load across all ages (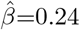, Table 2; nearly parallel loess curves, Figure 3). The same relationship was observed for diagnosed diabetics compared to non-diabetics, with 1.35-fold greater total load across all ages for diabetic participants.

When considering the effects of hypercholesterolemia (medicated for high cholesterol) and of high WHR ratio on log(WMH load), the loess-fitted curves suggested there might be a risk factor by age interaction (Figure 3). For both risk factors, the log(WMH load) seemed to be the same regardless of the risk factor status over the age of 70. Multiple linear regression was used to assess the importance of hypercholesterolemia by age interaction term. A new explanatory variable (multiple of the binary hypercholesterolemia variable and age) was added to model U.2.2, but there was not enough evidence to reject the null hypothesis of no effect (*p*=0.09). The WHR by age interaction was also explored (a new term added to model U.2.5) and it was marginally significant (*p*=0.049). However, the addition of the interaction term did not change *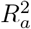* to two decimal places (no higher explanatory power was achieved). Thus both interaction terms were not further explored. Both hypercholesterolemia and WHR had marked positive effect on log(WMH load) (Table 2).

The highest risk groups for smoking and APOE-*ε* status (*>*50 pack years and APOE *ε*4/*ε*4) were associated with a higher log(WMH load) across all ages (Figure 3). In linear regression analyses, the effect of smoking and APOE-*ε* status on log(WMH load) was found to be strong and positive (Table 2). For example, comparing smoking score 1 to smoking score 0 (baseline) was associated with exp(0.15)=1.16-fold increase in WMH load, and 1.57-fold increase when comparing 2 to 0, respectively.

### 3.3. Spatial distribution of white matter hyperintensities

We plotted WMH incidence, voxel-wise, across 13,680 healthy ageing individuals and reveal the expected spatial distribution of WMHs. The highest probabilities were concentrated around the periventricular areas and in deep white matter regions (Figure 4).

**Figure 4:**
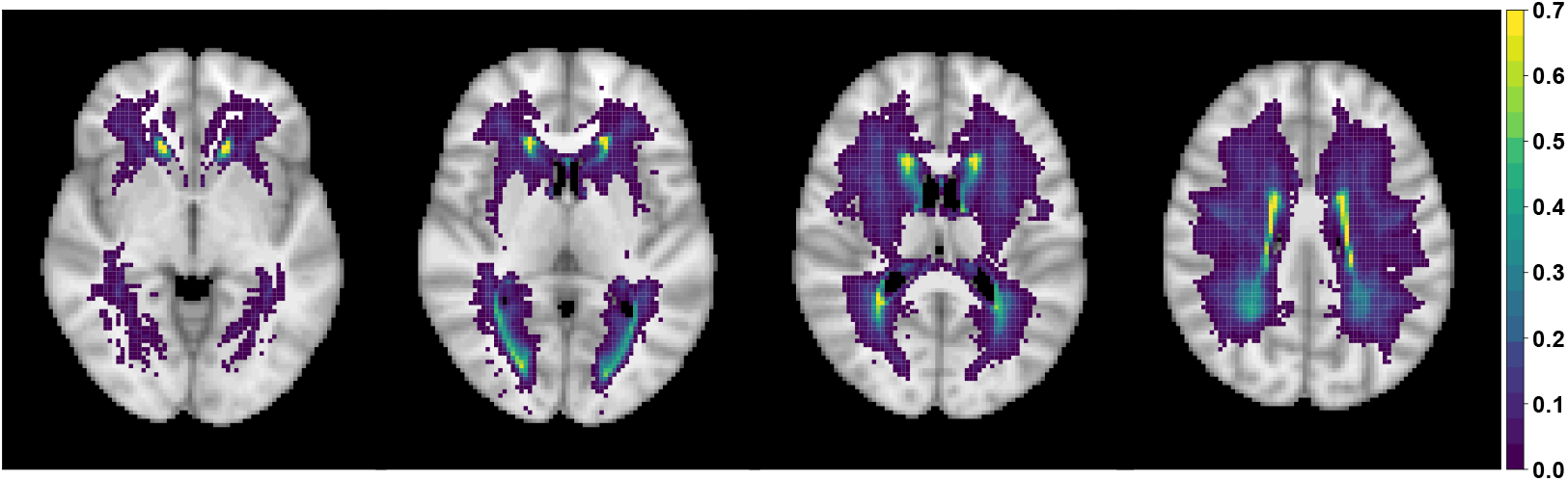
Square root transformed empirical WMH probability based on binary WMH masks of 13,680 UK Biobank individuals; axial slices *z*={35, 40, 45, 50} shown (from left to right). Square root transformation leads to more dispersed values allowing for better visualisation. Voxels with three or fewer individuals having a WMH are plotted as transparent to show a standard anatomical MRI for reference.

Next, we explored the effect of head size, age and sex, and their interaction on WMH probability (Figure 5) since they all act as confounding effects when considering the spatial distribution of WMHs associated with individual risk factors. The figure represents the standardized coefficients (z-scores based on mean BR estimates) for voxels which are significant (5% FDR correction applied) and have WMH incidence of at least 4 people (40,001 voxels, i.e. all other voxels are plotted as transparent). The age effect on WMH probability is dominant and widely spread through white matter. Also, note that the direction of the effect for those confounding variables is the same as in the univariate regression (Model U.1, Table 2), i.e. positive (red) for age and age by sex interaction, negative (blue) for sex and head size, but with varying effect size across the brain.

**Figure 5:**
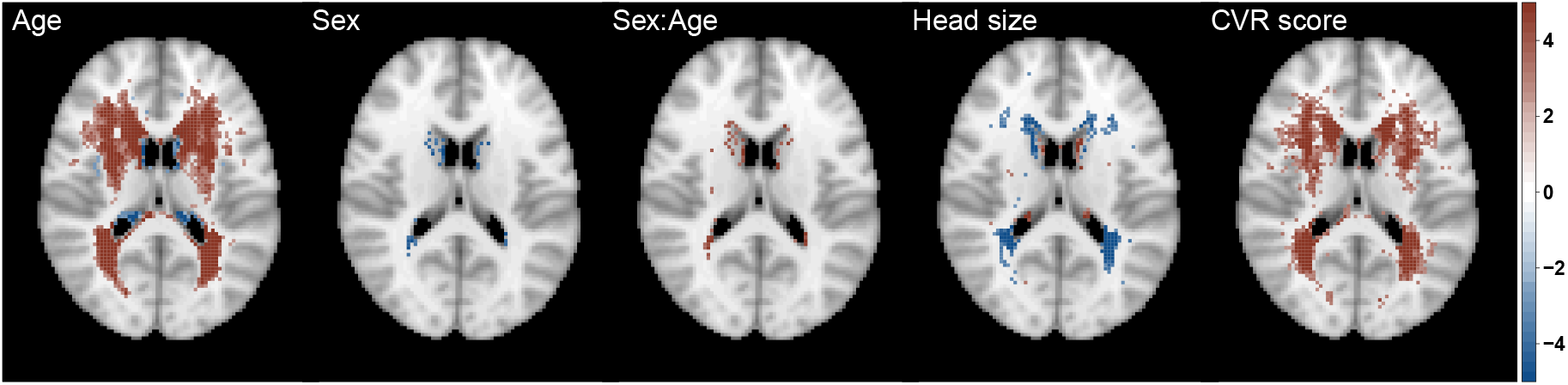
Significance maps (z-scores based on mean bias-reduced estimates) for model S.1, which includes age, sex (baseline men), age-sex interaction and head size and cerebrovascular risk (CVR) score as explanatory variables. Data on 13,680 UK Biobank individuals, and voxels with at least four individuals having a WMH explored (i.e. 0.03% WMH incidence); 5% FDR correction applied; axial slice *z*=45 shown.

We also explored the effect of each risk factor on WMH probability (marginal models, Figure 6(b)) as well as the contribution of each risk factor while controlling for all other risk factors (joint model, Figure 6(a)). On those axial slices, the darker the colour, the stronger the effect of the presence of the risk factor when compared to its absence. To quantify the WMHs associated with each risk factor, we estimated the percentage of significant voxels in a mask of 40,001 voxels for each risk factor in the marginal models and the change in the joint models, when other risk factors are controlled for (Table 3). This provided an additional measure of the relative importance of the different risk factors. The widest spatial distribution, in both periventricular and deep white matter, was for hypertension risk - thought to be the strongest risk factor, after age, for the presence of WMHs (Figure 6, Table 3).

**Figure 6:**
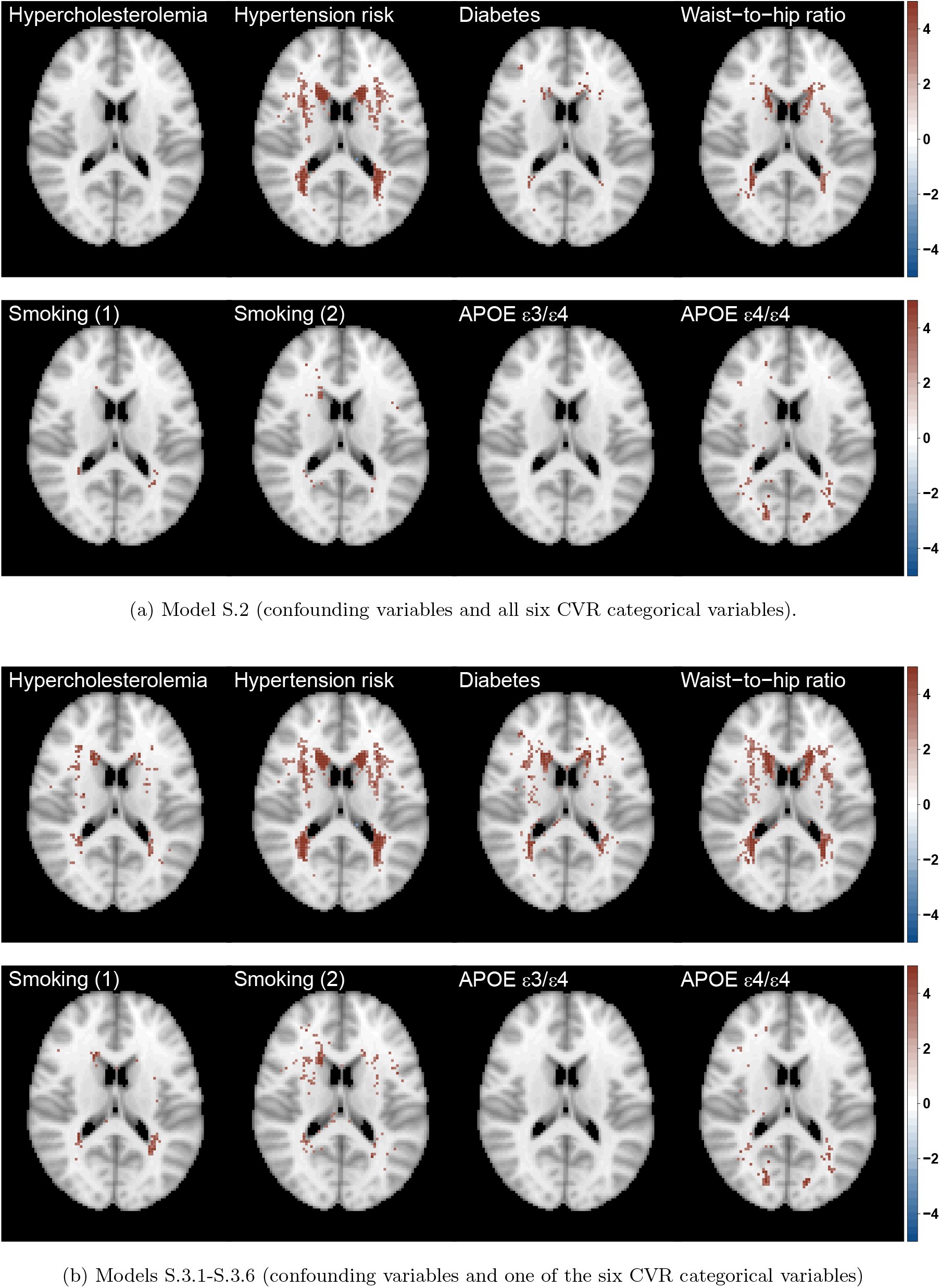
Significance maps (z-scores based on mean bias-reduced estimates) for (a) model S.2 (joint) and (b) models S.3.1 - S.3.6 (marginal). All models include the same confounding variables as models S.1 (age, sex (baseline men), age-sex interaction and head size). Data on 13,680 UK Biobank individuals, and voxels with at least four individuals having a WMH explored (i.e. 0.03% WMH incidence); 5% FDR correction applied; axial slice *z*=45 shown.

In terms of spatial extent, WHR and APOE *ε*4/*ε*4 genotype stand out as the next notable risk factors with a unique spatial distribution, independent of other risk factors. The contribution of APOE *ε*4/*ε*4 genotype persists regardless of the inclusion of the other risk factors in the model (significantly affecting about 4% of the voxels analysed, Table 3). We further plotted the APOE *ε*4/*ε*4 effect and found it was associated with WMH load concentrated at the boundary of the occipital and temporal lobes (Figure A.1). While the spatial distribution of WMHs associated with WHR was concentrated on the periventricular areas and had a similar distribution to diabetes positive status. Notably, when examining the unique contribution of the risk factors (Figure 6(a)), diabetes was much reduced in spatial extent (drop from 8.2% in the marginal model to 1.5% in the joint model), suggesting its contribution typically seen in the literature may be confounded by other CVR factors. Hypertension, APOE-*ε* status and WHR remained important risk factors, even after controlling for other CVRs (Figure 6(a), Table 3).

The estimated effect of the composite variable CVR score on WMH probability (Figure 5) reflected the combined but varied effect of its constituent risk factors (Figure 6(a)).

Additional analyses were run to ensure our WHR findings were not driven by associated hypertension, using continuous WHR and systolic BP (as opposed to binary indicators used in the main analysis). We investigated whether the spatial distribution of WMHs is independently affected by WHR and systolic BP. Figure A.2 shows the WHR effect is wider in terms of spatial extent with more spatially spread unique effects in deep white matter (Figure A.2(d)).

### 3.4. Cerebrovascular risk and speed of processing

Univariate linear regression estimated the relationship between log(WMH load) and speed of processing (reaction time in the ‘Snap’ task (Table 4)). Log(WMH load) was strongly associated with reaction time (*p*=0.009). The association between CVR score and reaction time did not appear to be significant after adjusting for confounding. A model including total CVR burden along with log(WMH load) showed only log(WMH load) to be a significant predictor of reaction time (*p*=0.011) but no explanatory power was gained (*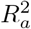* did not change).

**Table 4:**
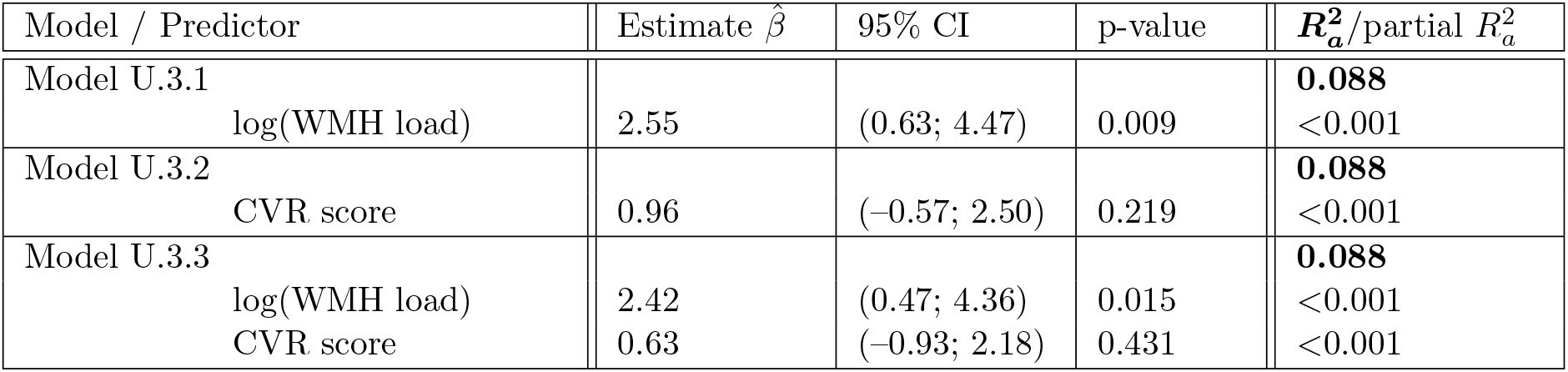
Univariate regression summaries outlining the association between the cognition variable reaction time (as outcome) and log(WMH load) and CVR score (as explanatory variables). Model U.3.1 - U.3.3: main effects of interest shown. All models adjusted for age, sex, head size, years of education and age-sex interaction. 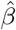 stands for the maximum likelihood estimate of the regression coefficient *β*.

Next, mediation analysis was used to determine whether any of the CV risks could explain the relationship between speed of processing and log(WMH load). Mediation analysis (Table 5) found WHR to be the only CVR factor to have a significant mediation effect (for the other risk factors, see Table A.4). Our results suggested 21% (9%; 83%) of the total effect of log(WMH load) on reaction time is explained through WHR.

**Table 5:** Mediation analysis. A significant proportion of the log(WMH load) (X) effect on speed of processing (Y) is explained through WHR mediation (mediator M). Models (mediator and outcome) are controlled for age, sex, age-by sex interaction, years of education and head size. See diagram A.3 for an illustration of the mediation analysis. 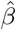 stands for the maximum likelihood estimate of the regression coefficient *β*.

| Average effect | WHR (continuous) |  |  |
| --- | --- | --- | --- |
| | Estimate $\hat{\beta}$ | 95% CI | p-value |
| $X$ on $M$ | 0.11 | (0.10; 0.13) | <0.001 |
| $M$ on $Y$ | 0.05 | (0.03; 0.08) | <0.001 |
| Mediation effect | 0.005 | (0.003; 0.010) | <10<sup>-15</sup> |
| Direct effect | 0.020 | (0.0003; 0.040) | 0.046 |
| Total effect | 0.025 | (0.006; 0.040) | 0.010 |
| Proportion mediated | 0.212 | (0.089; 0.830) | 0.010 |

## 4. Discussion

### White matter hyperintensity associated with individual risk factors

Population level brain imaging, lifestyle and demographic data on 13,680 healthy ageing volunteers were used to systematically investigate the association of cerebrovascular risk factors with the total burden and voxel-wise spatial distribution of WMHs. Contrary to previous reports, which typically emphasise hypertension as the main risk factor associated with WMH load, other cerebrovascular risk factors, including high waist-to-hip ratio, had similar or higher magnitude association with WMH burden, a unique spatial distribution and an independent relationship with cognition.

The contribution of known cerebrovascular risk factors to total WMH burden was examined. All CVR factors were found to be significant predictors of WMH load, motivating our exploration of the spatial distribution of the individual risk factors. Previous work in a subset (*N* =9, 722) of the UK Biobank cohort has shown independent contributions of hypertension, diabetes, WHR and smoking pack years to the total WMH load (Cox et al., 2019). We replicate this finding, showing an additional risk of homozygous APOE *ε*4 status and hypercholesterolemia in a larger cohort with complete data. Hypertension is frequently found to be the most predictive risk factor for total WMH load and therefore management of blood pressure is recommended to reduce both the total burden and the progression of WMHs (Verhaaren et al., 2013). There was a higher main effect size associated with the presence of diabetes compared to hypertension with the former associated with a 1.35-fold increase in total WMH burden compared to 1.27-fold increase for hypertension. However, risk factors including heavy smoking, APOE *ε*4/*ε*4 status and WHR had associations of similar magnitude, or greater, than hypertension. We found the presence of risk factors associated with higher total WMH load across all ages from 45–80 (Debette et al., 2011; Knopman et al., 2001). Together this points to the need for careful management of multiple risk factors across ageing for the preservation of brain vascular health.

A descriptive profile of this ageing population revealed overall higher cerebrovascular risk score (i.e. sum of risk factors) in males than females from mid through to late life. In terms of total WMH load, males and females had similar levels that increased linearly until age 65, after which total WMH load is higher in females. Several epidemiological studies corroborate this sex effect, with women appearing to consistently have a higher total WMH load than men (Sachdev et al., 2009). The age at which the difference in total WMH diverges between males and females has varied across studies, likely the result of limited sample sizes and age ranges within cohorts. One of the largest studies of WMH prevalence, the Rotterdam sample of healthy adults aged 60–90 (De Leeuw et al., 2001) found no sex differences in total WMH load, but did find sex differences across all decades in frontal periventricular WMH load (De Leeuw et al., 2001). As well as lower resolution scans (1.5 T) and a lower population size (*N* =1, 077) compared to ours (3T and *N* =13, 680), the study used a qualitative rating scale based on anatomical landmarks. Here, we used an objective method and voxel-wise test statistics across the whole brain to show sex effects and a sex by age interaction in 13,680 individuals concentrated in periventricular regions, with increased load in females over 65. This finding is important, given the higher risk of dementia associated with WMH burden and the higher prevalence of dementia in women. Notably, total CVR is higher in men of all ages, suggesting the increased WMH load seen in women aged over 65 may not be driven by cerebrovascular risk factors (or at least not the dominant risk factors included in our study). There is some evidence of a genetic component to WMHs, with higher heritability in women (Atwood et al., 2004; DeCarli et al., 1999). Another prominent hypothesis for higher total WMH load in women suggests the influence of sex hormones, particularly around menopause may be important. It is not yet known whether the higher incidence of AD in women is a risk or result of increased WMH load. Longitudinal data in UK Biobank (both imaging data and health outcomes) may help to establish if the increased WMH load in older women is associated with higher incidence of dementia.

### The spatial distribution of individual cerebrovascular risk factors

To date, the majority of studies have examined either total WMH load or deep versus periventricular WMHs associated with cerebrovascular risk factors. Where voxel-wise analyses have been attempted, this has either been using logistic regression (Lampe et al., 2019b), which we have discussed produces unstable test statistics or other “ad hoc” methods (Lampe et al., 2019a), but have not examined independent cerebrovascular risk factors voxel-wise.

Examination of the spatial distribution of individual cerebrovascular risk factors, led to several important findings. There were unique spatial patterns for certain risk factors and it was possible to quantify the contribution of different risk factors to total WMH load. Hypertension, WHR and APOE *ε*4 homozygosity emerge as the dominant risk factors in terms of the spatial extent of the probability of WMHs and the number of significant voxels after controlling for head size, age, sex and their interaction. However, both hypercholesterolemia and diabetes did not reveal consistent patterns of spatial distribution in models controlling for the other risk factors, despite the latter being an important predictor of total WMH load. The finding suggests these risk factors may interact with other risk factors such as hypertension and do not present a direct path to the pathogenesis of WMHs.

The homozygous *ε*4 genotype was revealed to be a significant predictor of WMH load. The finding is consistent with evidence of increased WMH volume in *ε*4 carriers in the UK Biobank cohort and longitudinal evidence of WMH progression associated with the *ε*4 genotype (Cox et al., 2017). Here we also replicate the finding from Lyall et al. (2019), in which there is no age interaction observed with APOE *ε*4 status in terms of total WMH load, contrary to some reports that APOE *ε*4 effects are most prominent in older age (Schiepers et al., 2012). Importantly, we extend these findings to show the spatial distribution of WMHs uniquely associated with the APOE *ε*4/*ε*4 genotype is concentrated in posterior deep white matter around the intracalcarine sulcus and extending superiorly into the temporal lobes. As evidence of the utility of our method, these independent effects of APOE *ε*4/*ε*4 status associated with WMHs in the deep white matter of the temporal-occipital lobes would not have been obvious with an approach examining only periventricular versus deep WMHs. The proximity of these WMHs to the medial temporal lobe is notable, given the susceptibility of this region to atrophy and disruption in AD. Given the association between APOE *ε*4 and dementia risk, it raises the question as to the potential contribution of WMHs to this risk. Longitudinal data is required to understand whether these WMHs have a role in the development or increased risk of dementia. Future studies would benefit from applying this voxel-wise method to examine how medial temporal lobe networks are impacted in the presence of WMHs in this region.

Waist-to-hip ratio above a healthy, sex-specific threshold, emerged as a critical risk factor for management in ageing. There was an independent spatial distribution of WMHs associated with WHR, that was not explained by comorbid hypertension, and was concentrated in the deep white matter and the ventricular caps. WHR was also the only risk factor to show a mediation effect on the relationship between speed of processing and total WMH burden. Waist circumference has been shown to be a reliable surrogate of visceral adiposity (Onat et al., 2004), and WHR is an especially useful measure in an ageing population because intra-abdominal fat tends to increase with age, whereas subcutaneous fat increases with degree of obesity but not age (Seidell et al., 1988). Cox et al. (2019), also noted an independent contribution of waist-to-hip ratio to WMH volume and suggest there may be metabolic and endocrine contributions to the pathogenesis of these WMHs that is distinct from arterial stiffness associated with high body mass (Cox et al., 2019). The dominance of WMHs associated with waist-to-hip ratio in deep white matter points to a possible ischaemic pathogenesis. Previous work examining deep versus periventricular WMHs associated with visceral obesity also found increased probability of WMHs in deep white matter associated with raised levels of proinflammatory cytokines (Lampe et al., 2019b). Proinflammatory cytokines are elevated in obesity and have been associated with cognitive decline and neurodegenerative processes associated with dementia (Pasha et al., 2017). This may help to explain the observed relationship to speed of processing we found with mediation analysis.

Here, we introduce a voxel-wise method that enables plotting the probability of the presence of WMHs associated with different risk factors or variables such as cognitive scores or symptoms such as depression or anxiety. The existing literature provides a confusing picture relating different cerebrovascular risk factors to the location of WMHs, and this is largely due to different classifications of lesion locations, either total load, separated into periventricular versus deep WMHs or divided between lobes or tracts. Our method provides a quantitative approach that can be used to standardise how WMHs are spatially identified across the brain. Our method has several applications, because of the granularity of spatial localisation that can be produced at a voxel-wise level. For example, future studies might integrate voxelwise results with other imaging modalities to examine how structural or functional networks are impacted by WMHs in specific regions.

### The relationship between risk factors and cognition

Reductions in speed of processing have been most frequently associated with total WMH load, but it is not clear if all WMHs contribute to this impairment or whether particular risk factors are implicated. It is now widely accepted that cognition is reliant on distributed brain networks. The spatial distribution of WHMs may therefore directly impact particular brain networks resulting in the observed cognitive deficits. Different risk factors may increase the likelihood of WMH in certain regions and therefore deferentially impact cognition. The conflicting evidence in the literature, as to whether WMHs do, or do not, correlate with cognitive performance may be in part due to conflation of WMHs associated with different CVR factors which present different spatial profiles. Beyond the spatial distribution, the contribution of individual risk factors to cognitive decline, controlling for dominant risk factors like hypertension, may inform clinical management strategies. In our analysis, WHR in particular showed a mediating effect on the relationship between speed of processing and total WMH burden. Future studies might profitably examine the longitudinal relationship between visceral adiposity and cognition to assess its importance as an early marker of cognitive decline. It is important to note that the explanatory power of the cognition models was relatively low which explains the magnitude of the regression coefficients, e.g. a change of 1 in log(WMH load) would only increase reaction time by 2.8ms, which is a significant effect but potentially not clinically meaningful.

### Limitations

Our findings should be interpreted in light of some limitations in the data used and biases existing within our cohort. There are limits to the conclusions that can be drawn from cross-sectional data, but the benefit of the cohort is a very large population size with a wide age range. Longitudinal studies are often limited in the age range and number of individuals that can be feasibly sampled over time. We used outputs from the UK Biobank pipeline for the estimation of WMHs. This pipeline did not directly account for ventricular size. Ventricular size is known to increase with age and so may affect segmentation of periventricular WMHs in older individuals. Nevertheless, the UK Biobank pipeline has a number of quality assurance steps to exclude individuals with large structural deviations (such as overly enlarged ventricles) and ensure accuracy of normalisation processes (Alfaro-Almagro et al., 2018). Due to sampling biases within the UK Biobank cohort (Fry et al., 2017) and our decision to exclude non-white individuals, our sample is limited in its generalisability to other ethnicities and sociodemographic groups. The UK Biobank sample is known to be generally healthier with higher socioeconomic status than the general UK population. To a certain extent, it is interesting to see effects of cerebrovascular risk factors in a relatively healthy cohort, and the effects in the general population may be much more pronounced. We were somewhat limited in the measurements we could use to represent different risk factors, with the majority being categorical variables. Continuous measure may have shown increased sensitivity. For example, WHR is a reliable surrogate marker of visceral adiposity (Onat et al., 2004) that is sensitive to age and metabolic syndromes (Seidell et al., 1988; Shuster et al., 2011), however it lacks the precision of CT or MRI for highly specific and comprehensive assessment of intra-abdominal fat. Future studies should assess intra-abdominal directly with CT or MRI. UK Biobank has collected such data but it was unavailable at the time of analysis.

### Conclusion

Our findings have some important clinical implications that may impact the management of cerebrovascular risk factors. We show the relative importance of different cerebrovascular risk factors in the contribution to total white matter hyperintensity burden in healthy ageing and the varied spatial distribution of CVR-related WMHs across the brain. Contrary to the assumption of hypertension as the dominant risk factor associated with WMH load, we show the associations of similar magnitude with APOE *ε*4/*ε*4 status, WHR, diabetes and heavy smoking. Independent and unique spatial distributions of WMHs associated with high WHR and APOE *ε*4/*ε*4 status point to careful management and observation of these risk factors.

Waist-to-hip ratio above healthy, sex-specific thresholds emerged as a key risk factor associated with WMHs in deep white matter and ventricular caps. There was some evidence of cognitive consequences to WHR-associated WMHs suggesting visceral adiposity, as indexed by WHR, represents a risk factor for close clinical management for mitigation of cognitive decline in healthy ageing.

## Data availability

The spatial distribution maps produced as part of the analysis are available at NeuroVault: https://neurovault.org/collections/AZQTNVUF/.

## Acknowledgements

Thank you to Dr Xin-You Tai for input on the design of the study. We would also like to thank Fidel Alfaro-Almagro, Ludovica Griffanti and Mark Jenkinson for their helpful advice regarding the UK Biobank imaging pipeline and BIANCA segmentation. Thank you to the participants of the UK Biobank for their time volunteering for the study.

## A. Complementary tables and figures

**Table A.1:**
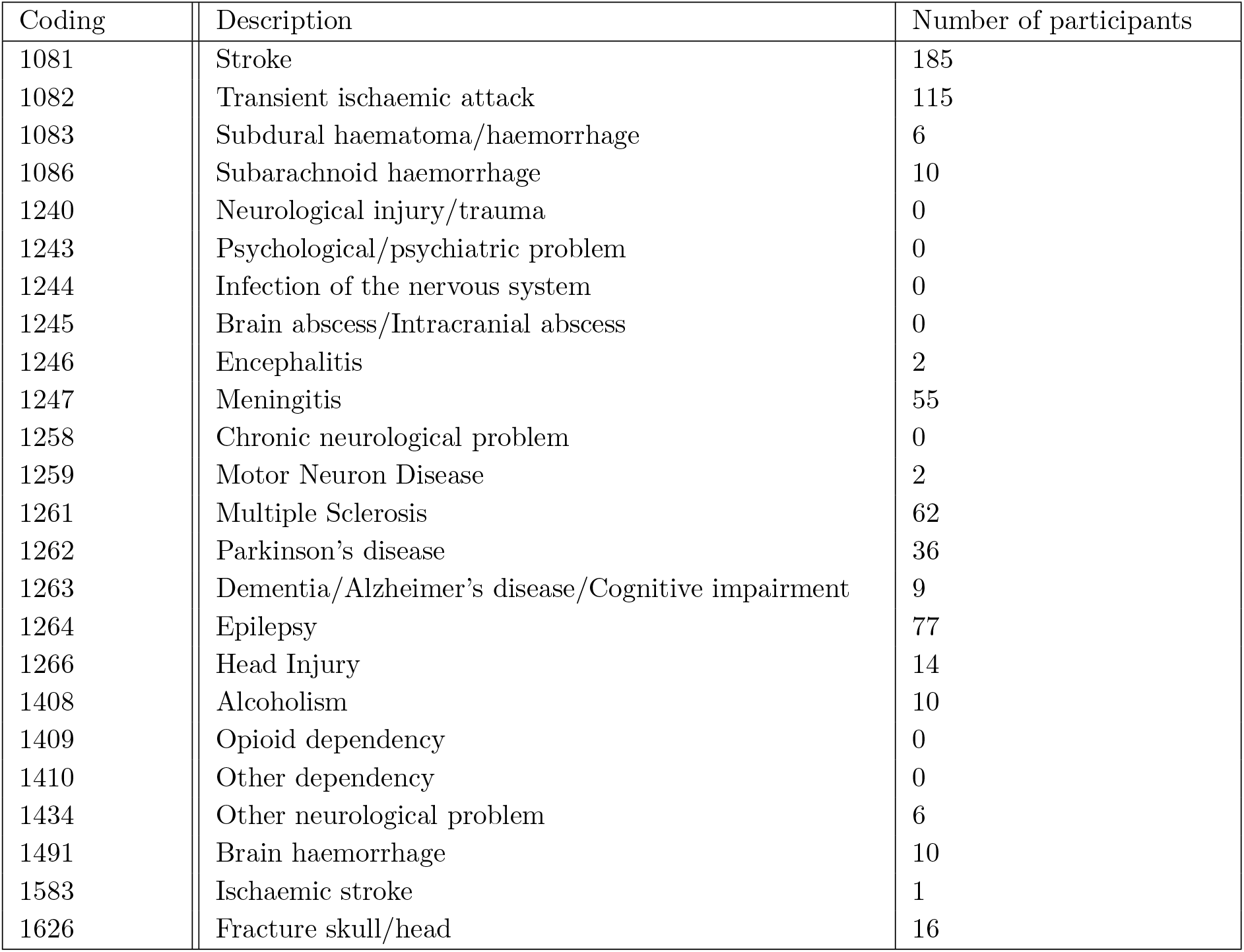
List of codes of the UK Biobank Data-field ‘Non-cancel illness’ (http://biobank.ndph.ox.ac.uk/showcase/field.cgi?id=20002) used for exclusion of participants in the data cleaning process. 616 illnesses reported across 590 participants.

**Table A.2:**
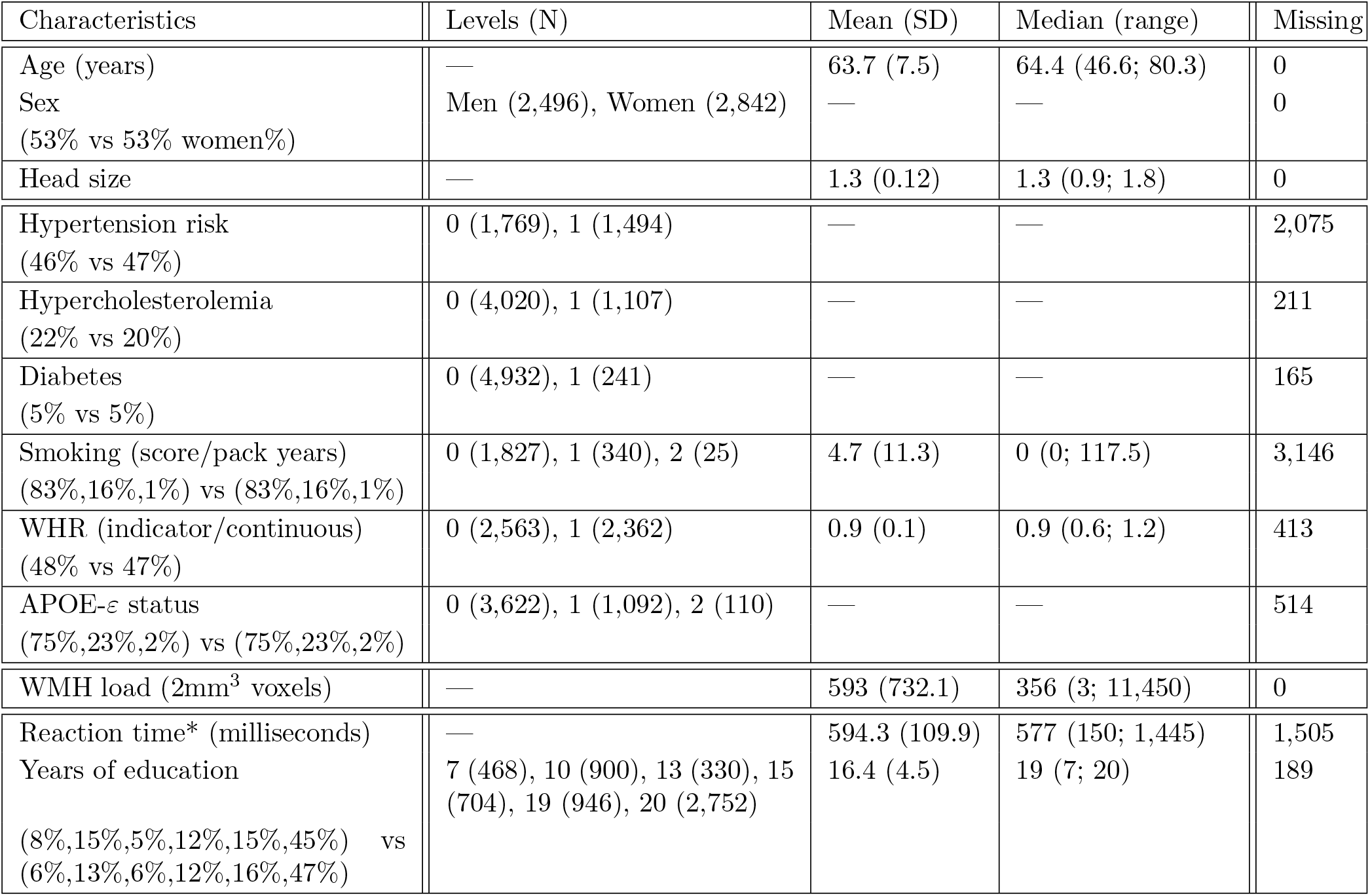
Characteristics of UK Biobank dataset on 5,338 participants, who were excluded from the analysis due to missingness (*6289 participants for the cognition variable reaction time and years of education). There does not seem to be systematic differences between the two datasets, so the exclusions are not likely to have introduced some bias in the results of the analyses. Similar to Table 1 with an additional column summarising the number of missing values for each variable. Summaries calculated without missing values for each variable. The percentage values under each discrete variable in column 1 are (% of participants with risk factor present (level 1) in exclusions dataset (calculated from Table A.2)) vs (% of participants with risk factor present in dataset used for the analyses (calculated from Table 1)), note for variables with more levels, vectors of incidences are presented. SD: standard deviation; WHR: waist-to-hip ratio; APOE: apolypoprotein-E; CVR: cerebrovascular risk; WMH: white matter hyperintensity.

| Characteristics | Levels (N) | Mean (SD) | Median (range) | Missing |
| --- | --- | --- | --- | --- |
| Age (years) | — | 63.7 (7.5) | 64.4 (46.6; 80.3) | 0 |
| Sex<br>(53% vs 53% women%) | Men (2,496), Women (2,842) | — | — | 0 |
| Head size | — | 1.3 (0.12) | 1.3 (0.9; 1.8) | 0 |
| Hypertension risk<br>(46% vs 47%) | 0 (1,769), 1 (1,494) | — | — | 2,075 |
| Hypercholesterolemia<br>(22% vs 20%) | 0 (4,020), 1 (1,107) | — | — | 211 |
| Diabetes<br>(5% vs 5%) | 0 (4,932), 1 (241) | — | — | 165 |
| Smoking (score/pack years)<br>(83%,16%,1%) vs (83%,16%,1%) | 0 (1,827), 1 (340), 2 (25) | 4.7 (11.3) | 0 (0; 117.5) | 3,146 |
| WHR (indicator/continuous)<br>(48% vs 47%) | 0 (2,563), 1 (2,362) | 0.9 (0.1) | 0.9 (0.6; 1.2) | 413 |
| APOE-ε status<br>(75%,23%,2%) vs (75%,23%,2%) | 0 (3,622), 1 (1,092), 2 (110) | — | — | 514 |
| WMH load (2mm<sup>3</sup> voxels) | — | 593 (732.1) | 356 (3; 11,450) | 0 |
| Reaction time* (milliseconds) | — | 594.3 (109.9) | 577 (150; 1,445) | 1,505 |
| Years of education<br>(8%,15%,5%,12%,15%,45%) vs<br>(6%,13%,6%,12%,16%,47%) | 7 (468), 10 (900), 13 (330), 15 (704), 19 (946), 20 (2,752) | 16.4 (4.5) | 19 (7; 20) | 189 |

**Table A.3:**
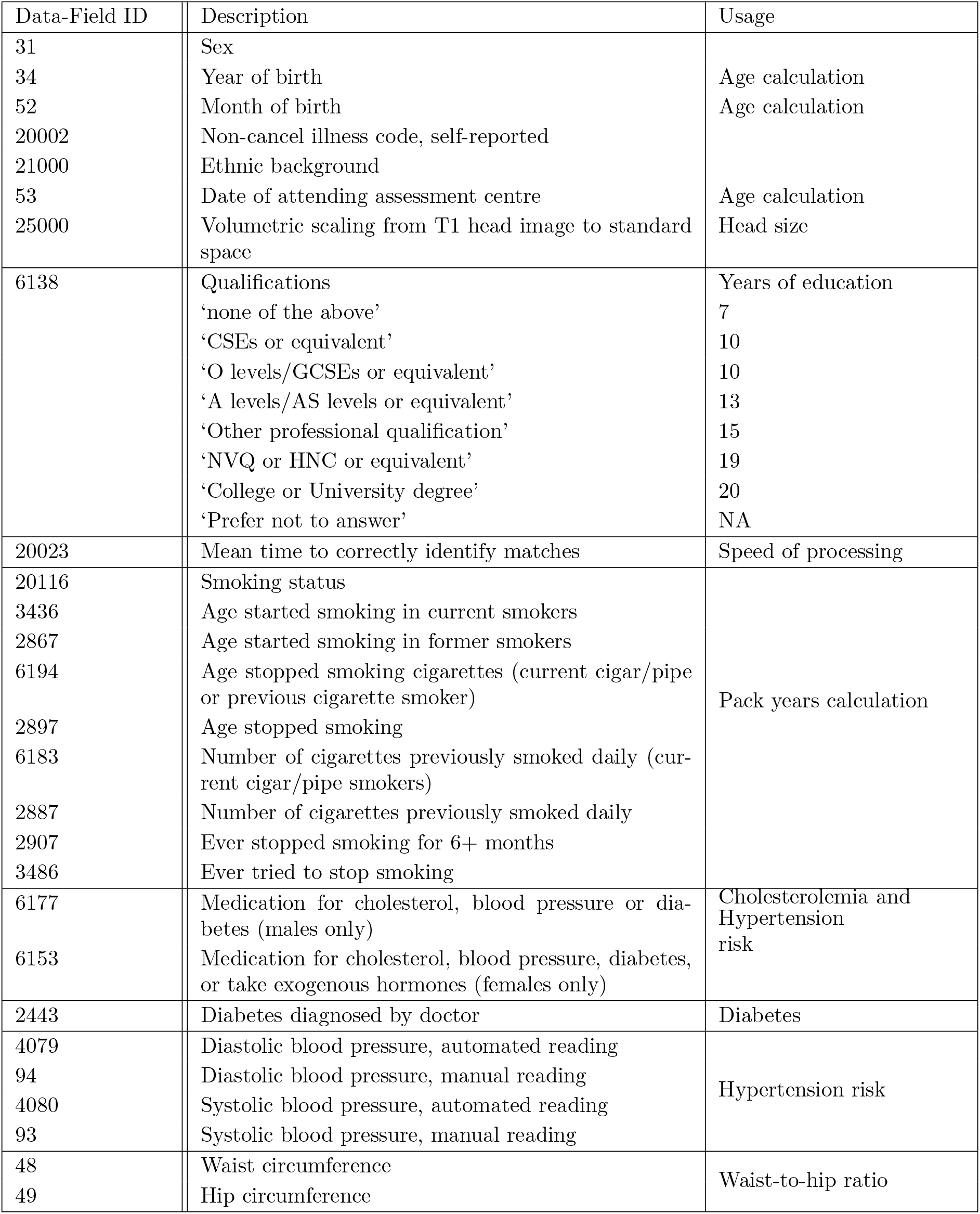
List of UK Biobank variables (available at http://biobank.ndph.ox.ac.uk/showcase/search.cgi used in the analysis. Data on APOE-*ε* status and WMH masks are not part of the catalogue.

**Figure A.1:**
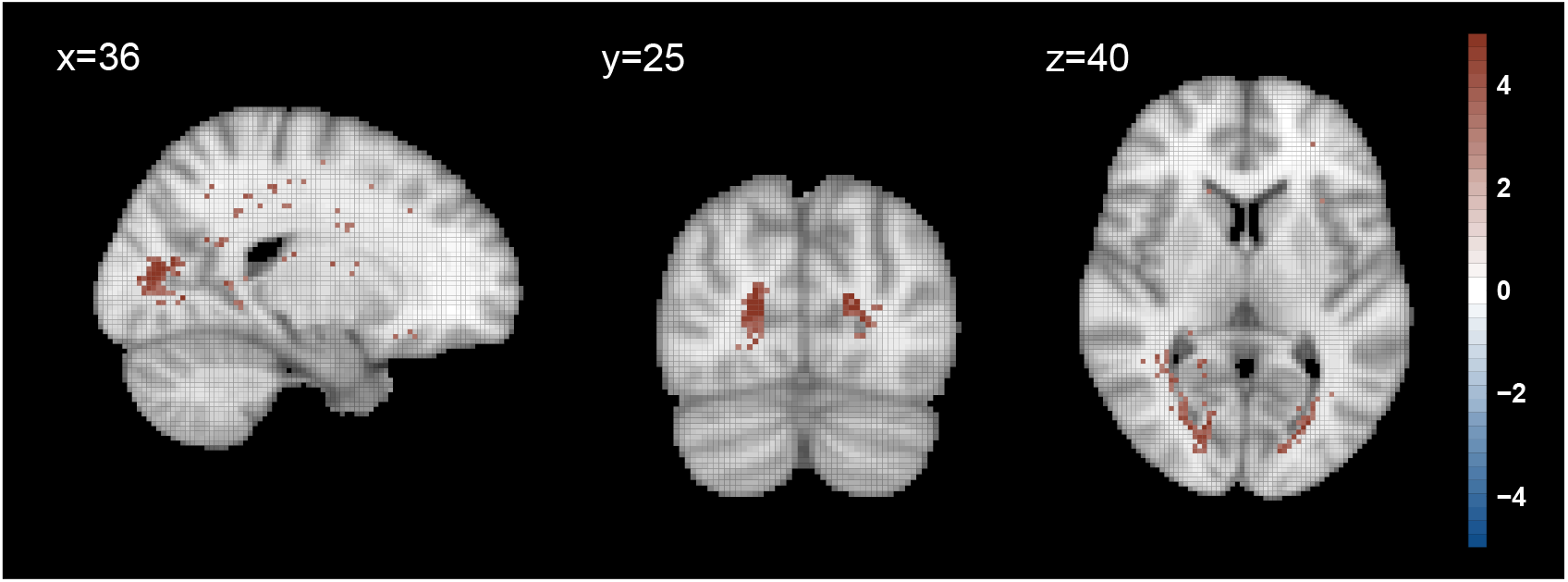
Significance maps (z-scores based on mean bias-reduced estimates) for APOE *ε*4/*ε*4 effect compared to no *ε*4 alleles (marginal model). Data on 13,680 UK Biobank individuals, and voxels with at least four individuals having a WMH explored (i.e. 0.03% WMH incidence); 5% FDR correction applied; Slices *x*=36, *y*=25, *z*=40 shown.

**Figure A.2:**
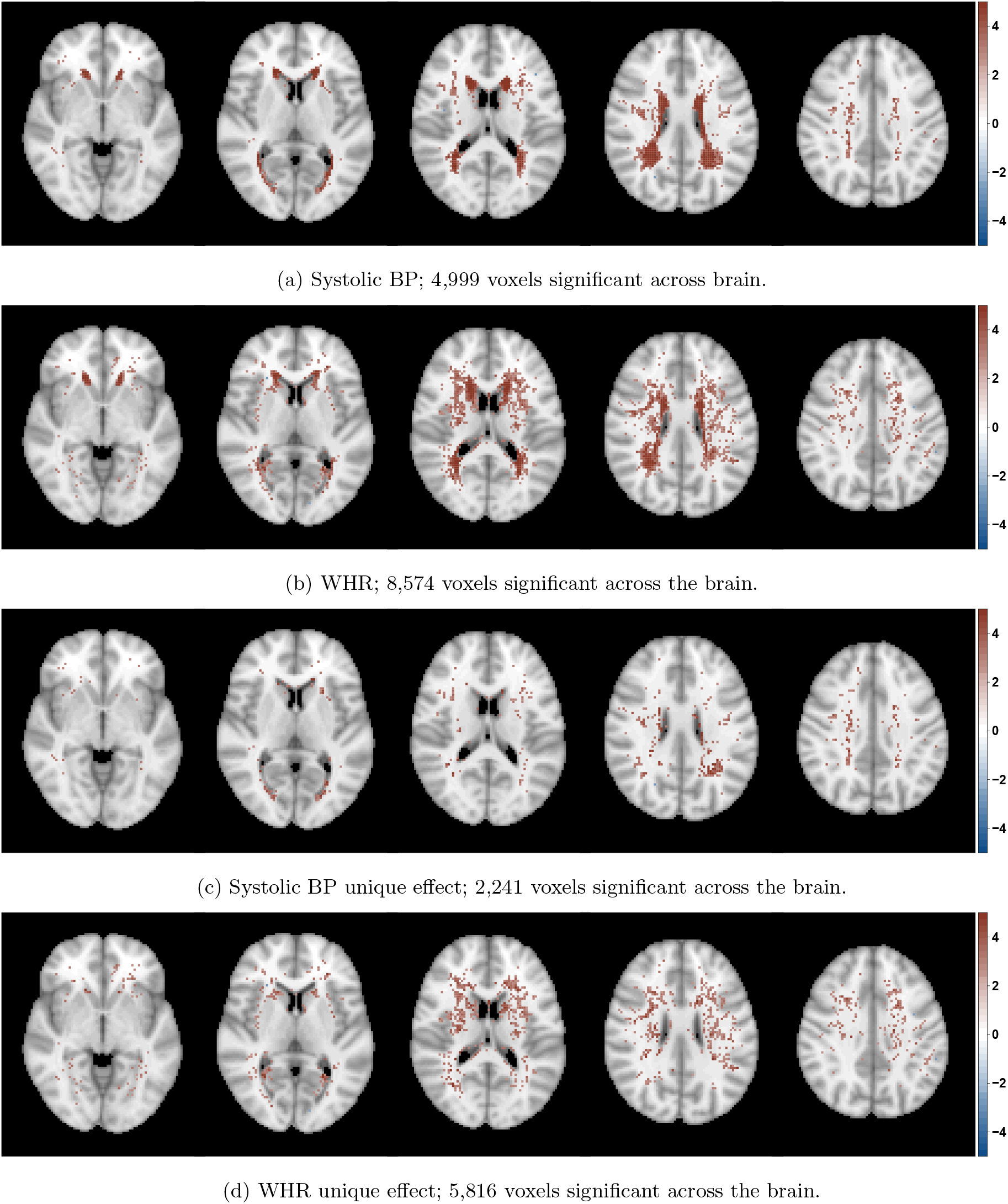
Significance maps (z-scores based on mean bias-reduced estimates) across five axial slices *z* = 35, 40, 45, 50, 55 in a model including age, sex, age-sex interaction, head size, systolic blood pressure and waist-to-hip ratio; data on 13,680 UK Biobank individuals and voxels with at least four individuals having a WMH explored; 5% FDR correction applied. From top to bottom each row shows z-scores for (a) Systolic BP, (b) WHR and their ‘unique’ effect, respectively, (c) ‘significant systolic BP/not significant WHR’ and (d) ‘not significant systolic BP/significant WHR’. BP: blood pressure; WHR: waist-to-hip ratio.

**Figure A.3:**
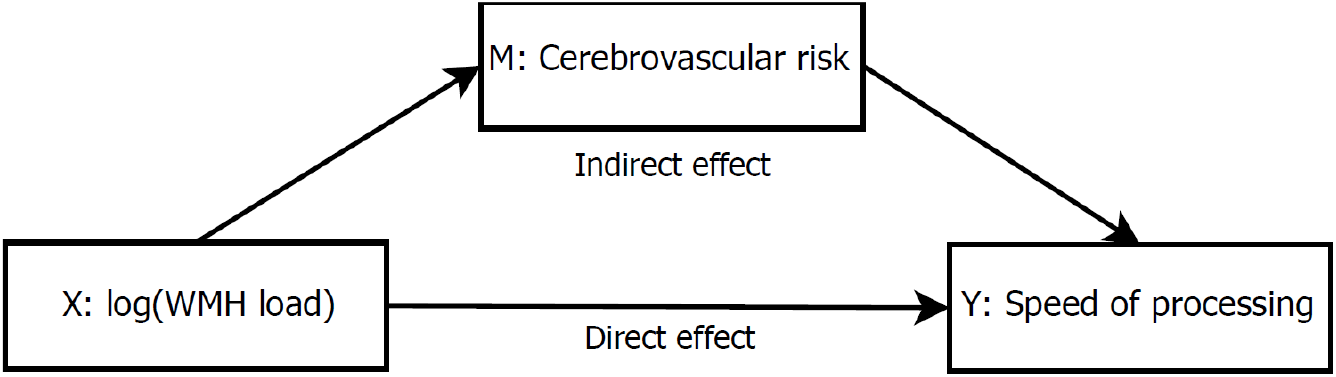
Mediation analysis diagram. The mediators explored are the six cerebrovascular risk factors. All models are controlled for age, sex, age-sex interaction, and head size. The main interest is in whether the indirect effect is significant, i.e. whether the effect of the predictor *X* on an outcome *Y* operates through a mediator variable *M* (fully or partially).

**Table A.4:** Mediation analysis exploring of the effect of log(WMH load) on speed of processing (reaction time) through cerebrovascular risk factors as mediator variable. Models (mediator and outcome) are controlled for age, sex, age-by sex interaction, years of education and head size. None of the risk factors shows a significant mediation effect. 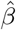 stands for the maximum likelihood estimate of the regression coefficient *β*. BP: blood pressure; CI: confidence interval. ^1^log-transformed pack years used (0.05 added to 0 pack year values); ^2^APOE-*ε* status represented by 1 if homozygous (*ε*4/*ε*4), 0 otherwise.

| Average effect | Systolic BP (continuous) |  |  | Hypertension risk (indicator) |  |  |
| --- | --- | --- | --- | --- | --- | --- |
| | Estimate $\hat{\beta}$ | 95% CI | p-value | Estimate $\hat{\beta}$ | 95% CI | p-value |
| Mediation effect | -0.001 | (-0.004; 0.000) | 0.342 | -0.002 | (-0.005; 0.00) | 0.222 |
| Direct effect | 0.027 | (0.008; 0.050) | 0.005 | 0.027 | (0.008; 0.050) | 0.005 |
| Total effect | 0.025 | (0.007; 0.040) | 0.006 | 0.025 | (0.006; 0.040) | 0.009 |
| Proportion mediated | -0.054 | (-0.296; 0.070) | 0.346 | -0.062 | (-0.346; 0.050) | 0.230 |

| Average effect | Hypercholesterolemia (indicator) |  |  | Diabetes (indicator) |  |  |
| --- | --- | --- | --- | --- | --- | --- |
| | Estimate $\hat{\beta}$ | 95% CI | p-value | Estimate $\hat{\beta}$ | 95% CI | p-value |
| Mediation effect | 0.001 | (-0.000; 0.000) | 0.141 | 0.001 | (-0.001; 0.000) | 0.227 |
| Direct effect | 0.024 | (0.005; 0.040) | 0.009 | 0.024 | (0.006; 0.040) | 0.012 |
| Total effect | 0.025 | (0.006; 0.040) | 0.007 | 0.025 | (0.007; 0.040) | 0.010 |
| Proportion mediated | 0.037 | (-0.016; 0.170) | 0.147 | 0.035 | (-0.027; 0.160) | 0.234 |

Table A.4: Mediation analysis exploring the effect of log(WMH load) on speed of processing (reaction time) through cerebrovascular risk factors as mediator variable.
| Average effect | Smoking<sup>1</sup> (continuous) | | | APOE-ε status (indicator<sup>2</sup>) | | |
| --- | --- | --- | --- | --- | --- | --- |
| | Estimate $\hat{\beta}$ | 95% CI | p-value | Estimate $\hat{\beta}$ | 95% CI | p-value |
| Mediation effect | -0.001 | (-0.002; 0.000) | 0.045 | 0.000 | (-0.001; 0.000) | 0.885 |
| Direct effect | 0.026 | (0.007; 0.050) | 0.005 | 0.025 | (0.007; 0.040) | 0.009 |
| Total effect | 0.025 | (0.006; 0.040) | 0.008 | 0.025 | (0.007; 0.040) | 0.010 |
| Proportion mediated | -0.043 | (-0.183; 0.000) | 0.053 | 0.002 | (-0.044; 0.060) | 0.886 |

## B. Justification of Minimum WMH Count

We chose a lower limit *Y*_min_ based on the following heuristic: At any one voxel, what is the smallest WMH count that can reject a null hypothesis of zero WMH incidence, *H*_0_ : *p* = 0? Of course, once a single WMH is observed we know *H*_0_ must be false, but as a heuristic it seems useful to assert that, if *fewer* than *Y*_min_ WHM are seen, we *cannot even* reject this obviously false *H*_0_, and this voxel should not be subject to further consideration.

This test takes the form 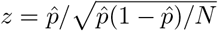, where 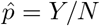. Solving *z ≥ z_α_* for *Y*_min_ shows that to obtain a minimum significance of *z_α_* requires 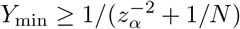. This result is virtually independent of *N*, giving *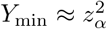*, and for *N* =13, 680 in particular, we found that *Y*_min_=4 was required to produce a *z* of at least 2, and hence we only considered voxels where 4 or more participants had a WMH.

## Footnotes

1 http://biobank.ctsu.ox.ac.uk/crystal/crystal/docs/brain_mri.pdf

2 https://fsl.fmrib.ox.ac.uk/fsl/fslwiki

3 While logit link is often used, probit and logit links give very similar results and we used probit link for comparability with other Bayesian probit-link models we investigate.

## References

Abraham, H. M. A., Wolfson, L., Moscufo, N., Guttmann, C. R., Kaplan, R. F., and White, W. B. (2016). Cardiovascular risk factors and small vessel disease of the brain: Blood pressure, white matter lesions, and functional decline in older persons.

Albert, A. and Anderson, J. A. (1984). On the existence of maximum likelihood estimates in logistic regression models. Biometrika.

Alfaro-Almagro, F., Jenkinson, M., Bangerter, N. K., Andersson, J. L., Griffanti, L., Douaud, G., Sotiropoulos, S. N., Jbabdi, S., Hernandez-Fernandez, M., Vallee, E., Vidaurre, D., Webster, M., McCarthy, P., Rorden, C., Daducci, A., Alexander, D. C., Zhang, H., Dragonu, I., Matthews, P. M., Miller, K. L., and Smith, S. M. (2018). Image processing and Quality Control for the first 10,000 brain imaging datasets from UK Biobank. NeuroImage.

Alfaro-Almagro, F., McCarthy, P., Afyouni, S., Andersson, J. L., Bastiani, M., Miller, K. L., Nichols, T. E., and Smith, S. M. (2020). Confound modelling in UK Biobank brain imaging. NeuroImage, page 117002.

Anderson, J., Jenkinson, M., and Smith, S. (2007). Technical report: Non-linear registration aka spatial normalisation. Technical report, FMRIB Centre, University of Oxford.

Andersson, J. L. R., Jenkinson, M., and Smith, S. M. (2007). Non-linear registration aka spatial normalisation. Internal Technical Report TR07JA1, Oxford Centre for Functional Magnetic Resonance Imaging of the Brain, Department of Clinical Neurology, Oxford University, Oxford, UK.

Atwood, L. D., Wolf, P. A., Heard-Costa, N. L., Massaro, J. M., Beiser, A., D’Agostino, R. B., and DeCarli, C. (2004). Genetic variation in white matter hyperintensity volume in the Framingham study. Stroke.

Benjamin, E. J., Blaha, M. J., Chiuve, S. E., Cushman, M., Das, S. R., Deo, R., De Ferranti, S. D., Floyd, J., Fornage, M., Gillespie, C., Isasi, C. R., Jim’nez, M. C., Jordan, L. C., Judd, S. E., Lackland, D., Lichtman, J. H., Lisabeth, L., Liu, S., Longenecker, C. T., MacKey, R. H., Matsushita, K., Mozaffarian, D., Mussolino, M. E., Nasir, K., Neumar, R. W., Palaniappan, L., Pandey, D. K., Thiagarajan, R. R., Reeves, M. J., Ritchey, M., Rodriguez, C. J., Roth, G. A., Rosamond, W. D., Sasson, C., Towfghi, A., Tsao, C. W., Turner, M. B., Virani, S. S., Voeks, J. H., Willey, J. Z., Wilkins, J. T., Wu, J. H., Alger, H. M., Wong, S. S., and Muntner, P. (2017). Heart Disease and Stroke Statistics’2017 Update: A Report from the American Heart Association.

Benjamini, Y. and Hochberg, Y. (1995). Controlling the False Discovery Rate: A Practical and Powerful Approach to Multiple Testing. Journal of the Royal Statistical Society: Series B (Methodological).

Boffa, R. J., Constanti, M., Floyd, C. N., and Wierzbicki, A. S. (2019). Hypertension in adults: Summary of updated NICE guidance.

Bycroft, C., Freeman, C., Petkova, D., Band, G., Elliott, L. T., Sharp, K., Motyer, A., Vukcevic, D., Delaneau, O., O’Connell, J., Cortes, A., Welsh, S., Young, A., Effingham, M., McVean, G., Leslie, S., Allen, N., Donnelly, P., and Marchini, J. (2018). The UK Biobank resource with deep phenotyping and genomic data. Nature.

Cheesman, R., Coleman, J., Rayner, C., Purves, K. L., Morneau-Vaillancourt, G., Glanville, K., Choi, S. W., Breen, G., and Eley, T. C. (2020). Familial Influences on Neuroticism and Education in the UK Biobank. Behavior Genetics, 50(2).

Cleveland, W. S., Grosse, E., and Shyu, W. M. (2017). Local regression models. In Statistical Models in S.

Cox, S. R., Lyall, D. M., Ritchie, S. J., Bastin, M. E., Harris, M. A., Buchanan, C. R., Fawns-Ritchie, C., Barbu, M. C., De Nooij, L., Reus, L. M., Alloza, C., Shen, X., Neilson, E., Alderson, H. L., Hunter, S., Liewald, D. C., Whalley, H. C., McIntosh, A. M., Lawrie, S. M., Pell, J. P., Tucker-Drob, E. M., Wardlaw, J. M., Gale, C. R., and Deary, I. J. (2019). Associations between vascular risk factors and brain MRI indices in UK Biobank. European Heart Journal.

Cox, S. R., Ritchie, S. J., Dickie, D. A., Pattie, A., Royle, N. A., Corley, J., Aribisala, B. S., Harris, S. E., Valdés Hernández, M., Gow, A. J., Muñoz Maniega, S., Starr, J. M., Bastin, M. E., Wardlaw, J. M., and Deary, I. J. (2017). Interaction of APOE e4 and poor glycemic control predicts white matter hyperintensity growth from 73 to 76. Neurobiology of Aging.

De Bresser, J., Kuijf, H. J., Zaanen, K., Viergever, M. A., Hendrikse, J., Biessels, G. J., Algra, A., Van Den Berg, E., Bouvy, W., Brundel, M., Heringa, S., Kappelle, L. J., Leemans, A., Luijten, P. R., Mali, W. P., Rutten, G. E., Vincken, K. L., and Zwanenburg, J. (2018). White matter hyperintensity shape and location feature analysis on brain MRI; Proof of principle study in patients with diabetes. Scientific Reports.

De Leeuw, F. E., De Groot, J. C., Achten, E., Oudkerk, M., Ramos, L. M., Heijboer, R., Hofman, A., Jolles, J., Van Gijn, J., and Breteler, M. M. (2001). Prevalence of cerebral white matter lesions in elderly people: A population based magnetic resonance imaging study. The Rotterdam Scan Study. Journal of Neurology Neurosurgery and Psychiatry, 70(1):9–14.

Debette, S. and Markus, H. S. (2010). The clinical importance of white matter hyperintensities on brain magnetic resonance imaging: Systematic review and meta-analysis.

Debette, S., Seshadri, S., Beiser, A., Au, R., Himali, J. J., Palumbo, C., Wolf, P. A., and DeCarli, C. (2011). Midlife vascular risk factor exposure accelerates structural brain aging and cognitive decline. Neurology, 77(5):461–468.

DeCarli, C., Reed, T., Miller, B. L., Wolf, P. A., Swan, G. E., and Carmelli, D. (1999). Impact of apolipoprotein E *ε*4 and vascular disease on brain morphology in men from the NHLBI twin study. Stroke.

Evans, D., Beckett, L., Albert, M., Hebert, L., Scherr, P., Funkenstein, H., and Taylor, J. (1993). Level of education and change in cognitive function in a community population of older persons. Annals of Epidemiology, 1(3):71–77.

Fawns-Ritchie, C. and Deary, I. J. (2020). Reliability and validity of the UK Biobank cognitive tests. PLoS ONE, 15(4).

Fazekas, F., Chawluk, J. B., and Alavi, A. (1987). MR signal abnormalities at 1.5 T in Alzheimer’s dementia and normal aging. American Journal of Neuroradiology.

Fazekas, F., Kleinert, R., Offenbacher, H., Schmidt, R., Kleinert, G., Payer, F., Radner, H., and Lechner, H. (1993). Pathologic correlates of incidental mri white matter signal hyperintensities. Neurology, 43(9):1683–1683.

Fazekas, F., Schmidt, R., and Scheltens, P. (1998). Pathophysiologic mechanisms in the development of age-related white matter changes of the brain. In Dementia and Geriatric Cognitive Disorders, volume 9.

Firth, D. (1993). Bias Reduction of Maximum Likelihood Estimates. Biometrika.

Fry, A., Littlejohns, T. J., Sudlow, C., Doherty, N., Adamska, L., Sprosen, T., Collins, R., and Allen, N. E. (2017). Comparison of Sociodemographic and Health-Related Characteristics of UK Biobank Participants with Those of the General Population. American Journal of Epidemiology.

Genovese, C. R., Lazar, N. A., and Nichols, T. (2002). Thresholding of statistical maps in functional neuroimaging using the false discovery rate. NeuroImage, 15(4).

Godin, O., Tzourio, C., Rouaud, O., Zhu, Y., Maillard, P., Pasquier, F., Crivello, F., Alpérovitch, A., Mazoyer, B., and Dufouil, C. (2010). Joint effect of white matter lesions and hippocampal volumes on severity of cognitive decline: The 3C-Dijon MRI study. Journal of Alzheimer’s Disease.

Green, P. J. (1984). Iteratively Reweighted Least Squares for Maximum Likelihood Estimation, and Some Robust and Resistant Alternatives. Journal of the Royal Statistical Society: Series B (Methodological).

Griffanti, L., Jenkinson, M., Suri, S., Zsoldos, E., Mahmood, A., Filippini, N., Sexton, C. E., Topiwala, A., Allan, C., Kivimäki, M., Singh-Manoux, A., Ebmeier, K. P., Mackay, C. E., and Zamboni, G. (2018). Classification and characterization of periventricular and deep white matter hyperintensities on MRI: A study in older adults.

Griffanti, L., Zamboni, G., Khan, A., Li, L., Bonifacio, G., Sundaresan, V., Schulz, U. G., Kuker, W., Battaglini, M., Rothwell, P. M., and Jenkinson, M. (2016). BIANCA (Brain Intensity AbNormality Classification Algorithm): A new tool for automated segmentation of white matter hyperintensities. NeuroImage.

Howard, V. J. (2013). Reasons underlying racial differences in stroke incidence and mortality. In Stroke.

Jeerakathil, T., Wolf, P. A., Beiser, A., Massaro, J., Seshadri, S., D’Agostino, R. B., and DeCarli, C. (2004). Stroke risk profile predicts white matter hyperintensity volume: The Framingham study. Stroke.

Jenkinson, M., Bannister, P., Brady, M., and Smith, S. (2002). Improved optimization for the robust and accurate linear registration and motion correction of brain images. NeuroImage, 17(2):825–841.

Kim, K. W., MacFall, J. R., and Payne, M. E. (2008). Classification of White Matter Lesions on Magnetic Resonance Imaging in Elderly Persons.

Kim, K. W., Seo, H., Kwak, M. S., and Kim, D. (2017). Visceral obesity is associated with white matter hyperintensity and lacunar infarct. International Journal of Obesity.

Knopman, D., Boland, L. L., Mosley, T., Howard, G., Liao, D., Szklo, M., McGovern, P., and Folsom, A. R. (2001). Cardiovascular risk factors and cognitive decline in middle-aged adults. Neurology, 56(1):42–48.

Kosmidis, I. (2020). brglm2: Bias Reduction in Generalized Linear Models. R package version 0.6.2.

Kosmidis, I. and Firth, D. (2009). Bias reduction in exponential family nonlinear models. Biometrika.

Kosmidis, I., Kenne Pagui, E. C., and Sartori, N. (2020). Mean and median bias reduction in generalized linear models. Statistics and Computing.

Lampe, L., Kharabian-Masouleh, S., Kynast, J., Arelin, K., Steele, C. J., Löffler, M., Witte, A. V., Schroeter, M. L., Villringer, A., and Bazin, P. L. (2019a). Lesion location matters: The relationships between white matter hyperintensities on cognition in the healthy elderly. Journal of Cerebral Blood Flow and Metabolism, 39(1):36–43.

Lampe, L., Zhang, R., Beyer, F., Huhn, S., Kharabian Masouleh, S., Preusser, S., Bazin, P. L., Schroeter, M. L., Villringer, A., and Witte, A. V. (2019b). Visceral obesity relates to deep white matter hyperintensities via inflammation. Annals of Neurology.

Lee, J. J., Wedow, R., Okbay, A., Kong, E., Maghzian, O., Zacher, M., Nguyen-Viet, T. A., Bowers, P., Sidorenko, J., Karlsson Linnér, R., Fontana, M. A., Kundu, T., Lee, C., Li, H., Li, R., Royer, R., Timshel, P. N., Walters, R. K., Willoughby, E. A., Yengo, L., Agee, M., Alipanahi, B., Auton, A., Bell, R. K., Bryc, K., Elson, S. L., Fontanillas, P., Hinds, D. A., McCreight, J. C., Huber, K. E., Litterman, N. K., McIntyre, M. H., Mountain, J. L., Noblin, E. S., Northover, C. A., Pitts, S. J., Sathirapongsasuti, J. F., Sazonova, O. V., Shelton, J. F., Shringarpure, S., Tian, C., Vacic, V., Wilson, C. H., Beauchamp, J. P., Pers, T. H., Rietveld, C. A., Turley, P., Chen, G. B., Emilsson, V., Meddens, S. F. W., Oskarsson, S., Pickrell, J. K., Thom, K., Timshel, P., Vlaming, R. d., Abdellaoui, A., Ahluwalia, T. S., Bacelis, J., Baumbach, C., Bjornsdottir, G., Brandsma, J. H., Concas, M. P., Derringer, J., Furlotte, N. A., Galesloot, T. E., Girotto, G., Gupta, R., Hall, L. M., Harris, S. E., Hofer, E., Horikoshi, M., Huffman, J. E., Kaasik, K., Kalafati, I. P., Karlsson, R., Kong, A., Lahti, J., van der Lee, S. J., Leeuw, C. d., Lind, P. A., Lindgren, K. O., Liu, T., Mangino, M., Marten, J., Mihailov, E., Miller, M. B., van der Most, P. J., Oldmeadow, C., Payton, A., Pervjakova, N., Peyrot, W. J., Qian, Y., Raitakari, O., Rueedi, R., Salvi, E., Schmidt, B., Schraut, K. E., Shi, J., Smith, A. V., Poot, R. A., St Pourcain, B., Teumer, A., Thorleifsson, G., Verweij, N., Vuckovic, D., Wellmann, J., Westra, H. J., Yang, J., Zhao, W., Zhu, Z., Alizadeh, B. Z., Amin, N., Bakshi, A., Baumeister, S. E., Biino, G., Bønnelykke, K., Boyle, P. A., Campbell, H., Cappuccio, F. P., Davies, G., De Neve, J. E., Deloukas, P., Demuth, I., Ding, J., Eibich, P., Eisele, L., Eklund, N., Evans, D. M., Faul, J. D., Feitosa, M. F., Forstner, A. J., Gandin, I., Gunnarsson, B., Halldórsson, B. V., Harris, T. B., Heath, A. C., Hocking, L. J., Holliday, E. G., Homuth, G., Horan, M. A., Hottenga, J. J., de Jager, P. L., Joshi, P. K., Jugessur, A., Kaakinen, M. A., Kähönen, M., Kanoni, S., Keltigangas-Järvinen, L., Kiemeney, L. A., Kolcic, I., Koskinen, S., Kraja, A. T., Kroh, M., Kutalik, Z., Latvala, A., Launer, L. J., Lebreton, M. P., Levinson, D. F., Lichtenstein, P., Lichtner, P., Liewald, D. C., Loukola, Life Lines Cohort Study, A., Madden, P. A., Mägi, R., Mäki-Opas, T., Marioni, R. E., Marques-Vidal, P., Meddens, G. A., McMahon, G., Meisinger, C., Meitinger, T., Milaneschi, Y., Milani, L., Montgomery, G. W., Myhre, R., Nelson, C. P., Nyholt, D. R., Ollier, W. E., Palotie, A., Paternoster, L., Pedersen, N. L., Petrovic, K. E., Porteous, D. J., Räikkönen, K., Ring, S. M., Robino, A., Rostapshova, O., Rudan, I., Rustichini, A., Salomaa, V., Sanders, A. R., Sarin, A. P., Schmidt, H., Scott, R. J., Smith, B. H., Smith, J. A., Staessen, J. A., Steinhagen-Thiessen, E., Strauch, K., Terracciano, A., Tobin, M. D., Ulivi, S., Vaccargiu, S., Quaye, L., van Rooij, F. J., Venturini, C., Vinkhuyzen, A. A., Völker, U., Völzke, H., Vonk, J. M., Vozzi, D., Waage, J., Ware, E. B., Willemsen, G., Attia, J. R., Bennett, D. A., Berger, K., Bertram, L., Bisgaard, H., Boomsma, D. I., Borecki, I. B., Bültmann, U., Chabris, C. F., Cucca, F., Cusi, D., Deary, I. J., Dedoussis, G. V., van Duijn, C. M., Eriksson, J. G., Franke, B., Franke, L., Gasparini, P., Gejman, P. V., Gieger, C., Grabe, H. J., Gratten, J., Groenen, P. J., Gudnason, V., van der Harst, P., Hayward, C., Hoffmann, W., Hyppönen, E., Iacono, W. G., Jacobsson, B., Järvelin, M. R., Jöckel, K. H., Kaprio, J., Kardia, S. L., Lehtimäki, T., Lehrer, S. F., Magnusson, P. K., Martin, N. G., McGue, M., Metspalu, A., Pendleton, N., Penninx, B. W., Perola, M., Pirastu, N., Pirastu, M., Polasek, O., Posthuma, D., Power, C., Province, M. A., Samani, N. J., Schlessinger, D., Schmidt, R., Sørensen, T. I., Spector, T. D., Stefansson, K., Thorsteinsdottir, U., Thurik, A. R., Timpson, N. J., Tiemeier, H., Tung, J. Y., Uitterlinden, A. G., Vitart, V., Vollenweider, P., Weir, D. R., Wilson, J. F., Wright, A. F., Conley, D. C., Krueger, R. F., Smith, G. D., Hofman, A., Laibson, D. I., Medland, S. E., Meyer, M. N., Yang, J., Johannesson, M., Visscher, P. M., Esko, T., Koellinger, P. D., Cesarini, D., Benjamin, D. J., Alver, M., Bao, Y., Clark, D. W., Day, F. R., Kemper, K. E., Kleinman, A., Langenberg, C., Trampush, J. W., Verma, S. S., Wu, Y., Lam, M., Zhao, J. H., Zheng, Z., Boardman, J. D., Freese, J., Harris, K. M., Herd, P., Kumari, M., Lencz, T., Luan, J., Malhotra, A. K., Ong, K. K., Perry, J. R., Ritchie, M. D., Smart, M. C., Wareham, N. J., Robinson, M. R., Watson, C., and Turley, P. (2018). Gene discovery and polygenic prediction from a genome-wide association study of educational attainment in 1.1 million individuals. Nature Genetics, 50(8).

Lloyd-Jones, D. M., Wilson, P. W., Larson, M. G., Beiser, A., Leip, E. P., D’Agostino, R. B., and Levy, D. (2004). Framingham risk score and prediction of lifetime risk for coronary heart disease. American Journal of Cardiology.

Lubin, J. H., Couper, D., Lutsey, P. L., Woodward, M., Yatsuya, H., and Huxley, R. R. (2016). Risk of cardiovascular disease from cumulative cigarette use and the impact of smoking intensity. Epidemiology.

Lyall, D., Cox, S., Lyall, L., Celis-Morales, C., Cullen, B., Mackay, D., Ward, J., Strawbridge, R., Mcintosh, A., Sattar, N., Smith, D., Cavanagh, J., Deary, I., and Pell, J. (2019). Association between apoe e4 and white matter hyperintensity volume, but not total brain volume or white matter integrity. Brain Imaging and Behavior, pages 1–9.

McCarron, M. O., Delong, D., and Alberts, M. J. (1999). APOE genotype as a risk factor for ischemic cerebrovascular disease: A meta-analysis. Neurology.

Moroni, F., Ammirati, E., Rocca, M. A., Filippi, M., Magnoni, M., and Camici, P. G. (2018). Cardiovascular disease and brain health: Focus on white matter hyperintensities.

Mortamais, M., Artero, S., and Ritchie, K. (2013). Cerebral white matter hyperintensities in the prediction of cognitive decline and incident dementia.

Onat, A., Avci, G. S., Barlan, M., Uyarel, H., Uzunlar, B., and Sansoy, V. (2004). Measures of abdominal obesity assessed for visceral adiposity and relation to coronary risk. International journal of obesity and related metabolic disorders : journal of the International Association for the Study of Obesity, 28:1018–25.

Pasha, E. P., Birdsill, A., Parker, P., Elmenshawy, A., Tanaka, H., and Haley, A. P. (2017). Visceral adiposity predicts subclinical white matter hyperintensities in middle-aged adults. Obesity Research and Clinical Practice.

Prins, N. D., Van Dijk, E. J., Den Heijer, T., Vermeer, S. E., Jolles, J., Koudstaal, P. J., Hofman, A., and Breteler, M. M. (2005). Cerebral small-vessel disease and decline in information processing speed, executive function and memory. Brain.

Rorden, C. and Karnath, H. O. (2004). Using human brain lesions to infer function: A relic from a past era in the fMRI age? Nature Reviews Neuroscience, 5(10).

Rostrup, E., Gouw, A. A., Vrenken, H., Van Straaten, E. C., Ropele, S., Pantoni, L., Inzitari, D., Barkhof, F., and Waldemar, G. (2012). The spatial distribution of age-related white matter changes as a function of vascular risk factors-Results from the LADIS study. NeuroImage.

Ryu, W. S., Woo, S. H., Schellingerhout, D., Chung, M. K., Kim, C. K., Jang, M. U., Park, K. J., Hong, K. S., Jeong, S. W., Na, J. Y., Cho, K. H., Kim, J. T., Kim, B. J., Han, M. K., Lee, J., Cha, J. K., Kim, D. H., Lee, S. J., Ko, Y., Cho, Y. J., Lee, B. C., Yu, K. H., Oh, M. S., Park, J. M., Kang, K., Lee, K. B., Park, T. H., Lee, J., Choi, H. K., Lee, K., Bae, H. J., and Kim, D. E. (2014). Grading and interpretation of white matter hyperintensities using statistical maps. Stroke.

Sachdev, P. S., Parslow, R., Wen, W., Anstey, K. J., and Easteal, S. (2009). Sex differences in the causes and consequences of white matter hyperintensities. Neurobiology of Aging.

Salvado, G., Brugulat-Serrat, A., Sudre, C. H., Grau-Rivera, O., Suarez-Calvet, M., Falcon, C., Fauria, K., Cardoso, M. J., Barkhof, F., Molinuevo, J. L., and Domingo Gispert, J. (2019). Spatial patterns of white matter hyperintensities associated with Alzheimer’s disease risk factors in a cognitively healthy middle-aged cohort. Alzheimer’s Research & Therapy.

Schiepers, O. J., Harris, S. E., Gow, A. J., Pattie, A., Brett, C. E., Starr, J. M., and Deary, I. J. (2012). APOE E4 status predicts age-related cognitive decline in the ninth decade: Longitudinal follow-up of the Lothian Birth Cohort 1921. Molecular Psychiatry, 17(3):315–324.

Seidell, J. C., Oosterlee, A., Deurenberg, P., Hautvast, J. G., and Ruijs, J. H. (1988). Abdominal fat depots measured with computed tomography: Effects of degree of obesity, sex, and age. European Journal of Clinical Nutrition, 42(9):805–815.

Shuster, A., Patlas, M., and Pinthus, J. (2011). The clinical importance of visceral adiposity: A critical review of methods for visceral adipose tissue analysis. The British journal of radiology, 85:1–10.

Smith, S. M. and Nichols, T. E. (2009). Threshold-free cluster enhancement: Addressing problems of smoothing, threshold dependence and localisation in cluster inference. NeuroImage.

Strassburger, T. L., Lee, H. C., Daly, E. M., Szczepanik, J., Krasuski, J. S., Mentis, M. J., Salerno, J. A., DeCarli, C., Schapiro, M. B., and Alexander, G. E. (1997). Interactive effects of age and hypertension on volumes of brain structures. Stroke.

Sudre, C. H., Cardoso, M. J., Frost, C., Barnes, J., Barkhof, F., Fox, N., and Ourselin, S. (2017). APOE *ϵ*4 status is associated with white matter hyperintensities volume accumulation rate independent of AD diagnosis. Neurobiology of Aging.

Tingley, D., Yamamoto, T., Hirose, K., Keele, L., and Imai, K. (2014). Mediation: R package for causal mediation analysis. Journal of Statistical Software, 59(5):1–38.

Van Dijk, E. J., Breteler, M. M., Schmidt, R., Berger, K., Nilsson, L. G., Oudkerk, M., Pajak, A., Sans, S., De Ridder, M., Dufouil, C., Fuhrer, R., Giampaoli, S., Launer, L. J., and Hofman, A. (2004). The association between blood pressure, hypertension, and cerebral white matter lesions: Cardiovascular determinants of dementia study. Hypertension.

Verhaaren, B. F., Vernooij, M. W., De Boer, R., Hofman, A., Niessen, W. J., Van Der Lugt, A., and Ikram, M. A. (2013). High blood pressure and cerebral white matter lesion progression in the general population. Hypertension.

Wardlaw, J. M., Smith, E. E., Biessels, G. J., Cordonnier, C., Fazekas, F., Frayne, R., Lindley, R. I., O’Brien, J. T., Barkhof, F., Benavente, O. R., Black, S. E., Brayne, C., Breteler, M., Chabriat, H., De-Carli, C., de Leeuw, F. E., Doubal, F., Duering, M., Fox, N. C., Greenberg, S., Hachinski, V., Kilimann, I., Mok, V., Oostenbrugge, R. v., Pantoni, L., Speck, O., Stephan, B. C., Teipel, S., Viswanathan, A., Werring, D., Chen, C., Smith, C., van Buchem, M., Norrving, B., Gorelick, P. B., and Dichgans, M. (2013). Neuroimaging standards for research into small vessel disease and its contribution to ageing and neurodegeneration.

Wardlaw, J. M., Valdés Hernández, M. C., and Muñoz-Maniega, S. (2015). What are white matter hyperintensities made of? Relevance to vascular cognitive impairment.

Whalley, L. J., Deary, I. J., Appleton, C. L., and Starr, J. M. (2004). Cognitive reserve and the neurobiology of cognitive aging.

Wiseman, R. M., Saxby, B. K., Burton, E. J., Barber, R., Ford, G. A., and O’Brien, J. T. (2004). Hippocampal atrophy, whole brain volume, and white matter lesions in older hypertensive subjects. Neurology.

World Health Organization (2008). Waist Circumference and Waist-Hip Ratio. Report of a WHO Expert Consultation.

